# Spinal cord homeostatic plasticity gates mechanical allodynia in chronic pain

**DOI:** 10.1101/2022.01.10.475704

**Authors:** Bing Cao, Gregory Scherrer, Lu Chen

## Abstract

Central sensitization caused by disinhibition of spinal cord circuits is a key mechanism of mechanical allodynia in neuropathic pain. Despite intense efforts, the molecular mechanisms that drive disinhibition and induce allodynia after peripheral nerve injury remain unclear. Using the spared-nerve injury (SNI) model of allodynia, we here demonstrate that SNI induces disinhibition of spinal nociceptive circuits by triggering homeostatic synaptic plasticity. Specifically, SNI-triggered homeostatic plasticity suppresses the inhibitory outputs of parvalbumin-positive (PV+) interneurons that form synapses on both primary afferent terminals and excitatory interneurons, causing hyperactivation of the nociceptive pathway. Using genetic manipulations, we identified the retinoic acid receptor RARα as the key mediator of the homeostatic synaptic plasticity underlying this synaptic disinhibition. Deletion of RARα in PV+ neurons blocked SNI-induced spinal disinhibition, central sensitization, and allodynia. Moreover, deletion of RARα in spinal PV+ neurons or application of an RARα antagonist in the spinal cord prevented the development of SNI-induced mechanical allodynia. Together, our results reveal a molecular mechanism of neuropathic pain whereby homeostatic plasticity causes the mis-direction of tactile information flow to ascending nociceptive pathways following peripheral nerve injury.

## Main text

Mechanical allodynia, a persistent pain condition induced by tissue or nerve injury in which an innocuous mechanical stimulus becomes painful, is a prevalent form of chronic pain affecting up to 10% of the population^1^. Although multiple locations in the CNS respond to peripheral nerve injury^2,3^, the key circuit responsible for transforming normal touch perception into mechanical allodynia is believed to reside within the spinal dorsal horn^4-7^. After peripheral nerve injury, both synaptic and non-synaptic changes increase the sensitivity of somatosensory neurons in the dorsal horn to tactile inputs, resulting in central sensitization^8^. Many mechanisms have been proposed to drive central sensitization that results in hyperalgesia and allodynia^9-16^. One such proposal, the central inhibitory gate theory, places local inhibitory interneurons at key gate-keeping positions of the dorsal horn circuit^9,17,18^. According to this theory, reduction in dorsal horn inhibitory tone drives the circuit level changes underlying hyperalgesia and allodynia^7,19-24^. Indeed, manipulations that inhibit or enhance GABAergic/glycineric outputs in central circuits are effective in promoting or alleviating neuropathic pain, respectively^7,25-29^. In contrast to the rich literature on circuit changes that underlie the central sensitization leading to allodynic pain, the molecular and cellular mechanisms that produce the change in inhibitory tone after peripheral nerve injury are unclear.

In the present study, we explored how dorsal horn synaptic plasticity drives neuropathic pain. Specifically, we investigated the hypothesis that peripheral nerve injury induces excessive homeostatic synaptic compensation, resulting in a weakened inhibitory output from the gate-keeping interneurons, which in turn overexcites nociceptive projection neurons and results in mechanical allodynia. We demonstrate that spared nerve injury (SNI) of the sciatic nerve significantly reduces the synaptic output of inhibitory dorsal horn PV+ interneurons onto both primary afferent terminals and local excitatory relay interneurons. Moreover, we report that this form of homeostatic plasticity requires the retinoic acid receptor RARα, a key mediator for homeostatic synaptic plasticity at CNS synapses^30-33^. Consistent with the notion that allodynia is due to RARα-dependent homeostatic plasticity, genetic deletion of RARα in dorsal horn PV+ neurons or pharmacological inhibition of RARα in the spinal dorsal horn prevented homeostatic downregulation of synaptic inhibition from PV+ neurons, and thereby blocked the development of SNI-induced mechanical allodynia and chronic pain.

### SNI-induced mechanical allodynia requires RARα in PV+ neurons

To investigate the functional contribution of RARα in PV+ neurons to chronic pain, we selectively deleted RARα in PV+ neurons by crossing a PV-specific Cre driver line with mice bearing a RARα conditional allele (PV-RARα conditional knockout mice). A Cre-dependent YFP reporter allele was also included to enable identification of PV+ cells. Expression fidelity of the PV-Cre driver line was verified in three CNS regions, ACC, S1, and dorsal horn, known to be involved in pain processing (Fig. 1a, b). Cre-dependent deletion of RARα was confirmed in spinal cord sections via *in situ* hybridization (ISH) (Fig. 1c-1d) and single-cell qRT-PCR (Fig. 1e). These results indicate a successful deletion of RARα in dorsal horn PV+ neurons in PV-RARα cKO mice.

**Fig. 1.**
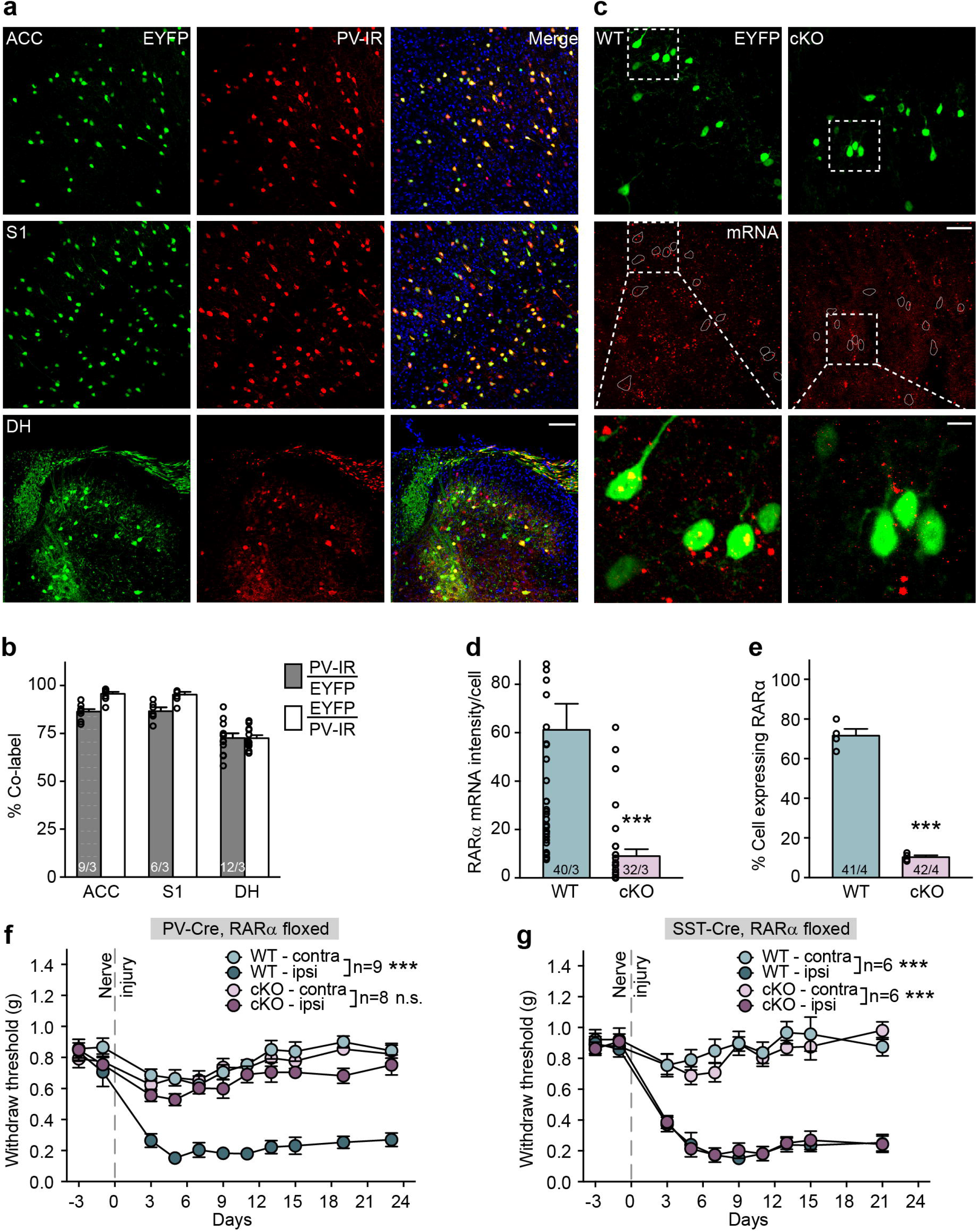
Deletion of RARα in parvalbumin-expressing (PV+) neurons blocks mechanical allodynia development. **a**. Representative images of anterior cingulate cortex (ACC), primary somatosensory cortex (S1) and spinal cord dorsal horn lumbar region (DH) showing Cre-dependent EYFP reporter (green) and PV-immunoreactivity (PV-IR, red) in PV-Cre::RARα cKO mice. Scale bar: 100 μm. **b**. Quantification of PV-Cre driver line expression fidelity (% PV-IR/EYFP) and efficacy (% EYFP/PV-IR) in ACC, S1 and DH. n/N = # of section/# of animals. **c**. Representative images of RNA scope in situ hybridization shows RARα mRNA expression (red) in spinal cord sections from WT and PV-RARα cKO mice. PV+ neurons are indicated by EYFP reporter (green). Scale bar: 50 μm. The overlay images of insets (square with white dashed line) are shown at bottom (scale bar: 10 μm). **d**. Quantification of RARα mRNA expression in WT (n/N = 40 cells/3 mice) and cKO mice (n/N = 32 cells/3 mice). Two-tailed unpaired t test: t = 4.2, ***, *p* < 0.001. **e**. Quantification of RARα mRNA expression percentage with single-cell qRT-PCR showing percent of RARα+ neurons in PV-expressing spinal dorsal horn neurons (WT: n/N = 41 cells/4 mice; cKO: n/N = 42 cells/4 mice). Two-tailed unpaired t test: t = 17.4, ***, *p* < 0.001. **f**. Mechanical allodynia quantified as paw withdrawal threshold with Von Frey test from hindlimbs ipsilateral and contralateral to the SNI side in WT and PV-RARα cKO mice before and over a period of 3 weeks following SNI surgery. Two-way ANOVA with post-hoc Bonferroni test, ***, *p* < 0.001. **g**. Mechanical allodynia quantified as paw withdrawal threshold with Von Frey test from hindlimbs ipsilateral and contralateral to the SNI side in WT and SST-RARα cKO mice before and over a period of 3 weeks following SNI surgery. Two-way ANOVA with post-hoc Bonferroni test, ***, *p* < 0.001. All graphs represent mean ± SEM.

Given RA and RARα signaling is known to play a central role in homeostatic synaptic plasticity in the cortex^30,31,34^ and hippocampus^32,33,35^, we performed a general behavioral survey. PV-RARα cKO mice exhibited normal behavior in the Y-maze (Extended Data Fig. 1a, b), elevated plus maze (Extended Data Fig. 1c, d), and open field test (Extended Data Fig. 1e – 1h), indicating normal working memory, locomotion and anxiety. Given the essential role of RARα signaling in PV+ neurons in gating inhibitory synaptic output in cortical sensory deprivation-induced homeostatic plasticity^36^, we next asked whether deletion of RARα in PV-expressing neurons impacted the development of neuropathic pain. To induce neuropathic pain, we used the SNI model of the sciatic nerve established in rodents^37,38^ (Extended Data Fig. 2a-e). Mechanical allodynia was assessed with the von Frey filament test three days after the injury and retested every other day for a minimum of three weeks.

Mechanical allodynia developed in the ipsilateral side of wildtype (WT) mice after nerve injury, and lasted for the duration of the observation. Interestingly, development of mechanical allodynia was completely blocked in PV-RARα cKO mice (Fig. 1f) but remain intact in somatostatin (SST)-specific RARα cKO mice (Fig. 1g). Thermal hyperalgesia resulted from SNI remained intact in PV-RARα cKO mice (Extended Data Fig. 2f), indicating that PV-RARα expression is selectively involved in neural circuit modifications underlying SNI-induced mechanical but not thermal allodynia. Moreover, responses to local inflammation induced by subcutaneous injection of 1% formalin were comparable between WT and PV-RARα cKO mice (Extended Data Fig. 3), suggesting that the PV-RARα deletion did not affect acute nociception.

### SNI-induced nociceptive pathway activation is reduced in PV-RARα cKO mice

In sham lesioned animals (WT or PV-RARα cKO), no significant neural activation, evaluated with FOS expression, was observed in the ipsilateral spinal cord dorsal horn in response to a light brush (Fig. 2a, b). By contrast, significantly greater FOS expression levels were found in the ipsilateral dorsal horn after SNI compared to the contralateral unstimulated side, indicating heightened sensitivity to light brush stimulation after injury (Fig. 2a, b). Importantly, deletion of RARα in PV+ neurons significantly reduced this hypersensitivity (Fig. 2b), supporting our results that RARα expression in PV+ neurons is critical for the development of mechanical allodynia (Fig. 1f).

**Fig. 2.**
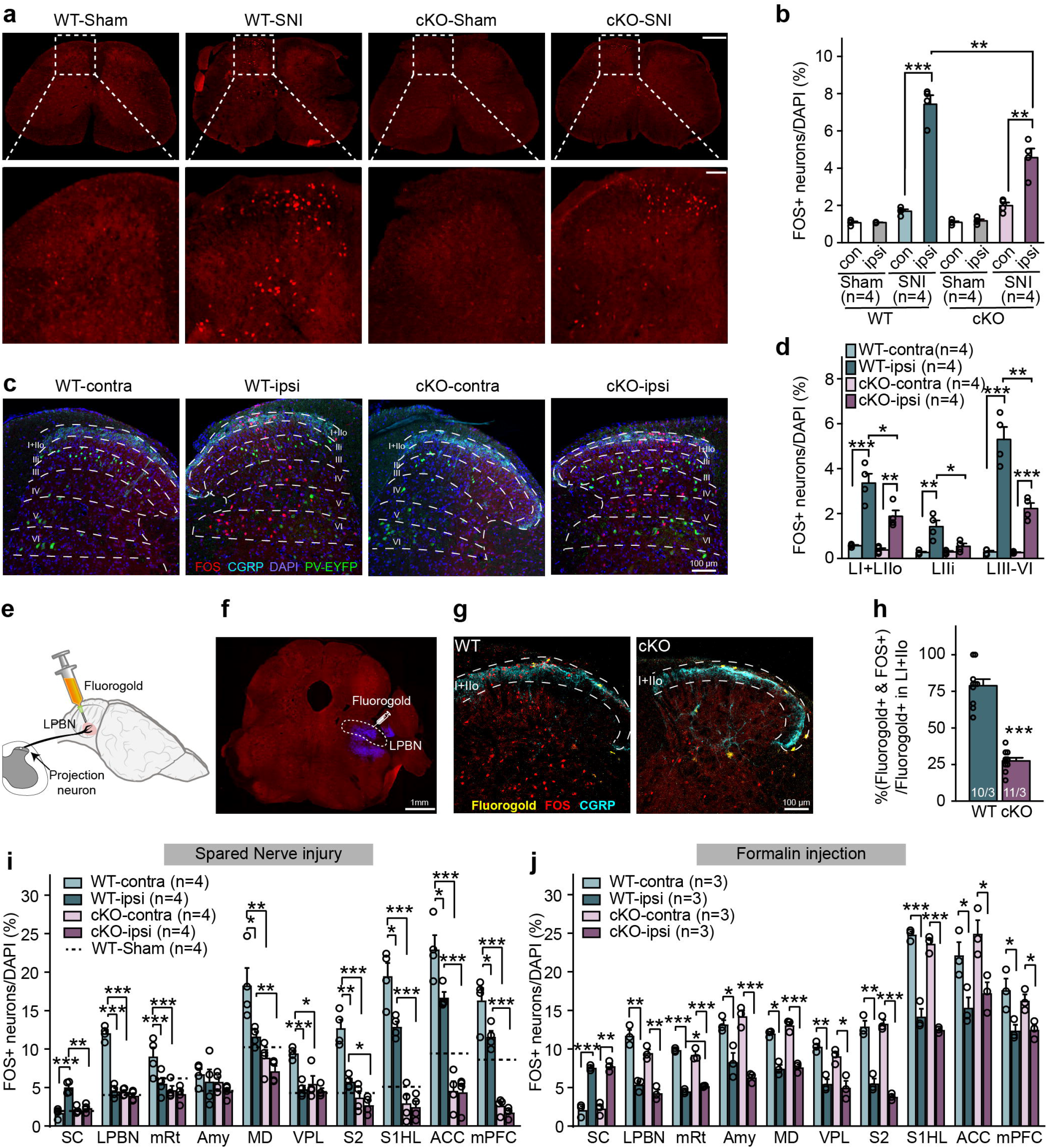
PV-RARα deletion reduces SNI-induced neuronal hyperactivity in the ascending nociceptive pathways. **a**. Representative images of spinal cord sections stained with FOS in WT and PV-RARα cKO mice after light brush stimulation. Images of ipsilateral dorsal horn were shown zoomed in at the bottom. Scale bars: 500 μm (top), 100 μm (bottom). **b**. Quantification of light brush-activated neurons (FOS+) in WT and PV-RARα cKO mice. **, *p* < 0.01; ***, *p* < 0.001. **c**. Representative images of dorsal horn sections stained with FOS (red), DAPI (blue), and CGRP (cyan, lamina I+IIo). PV+ cells are indicated by EYFP (Green). Scale bar: 100 μm. **d**. Quantification of neuronal activation in different dorsal horn laminae in WT and PV-RARα cKO mice with SNI after light brush stimulation. *, *p* < 0.05; **, *p* < 0.01; ***, *p* < 0.001. **e**. A schematic drawing depicting retrograde labeling of dorsal horn pain projection neurons via injection of fluorogold into LPBN, where the secondary order neurons receiving projections from dorsal horn are located. **f**. A representative image showing the fluorogold injection site (blue) in LPBN. Scale bar: 1 mm. **g**. Representative images showing retrogradely labeled projection neurons (yellow) in the dorsal horn superficial lamina (indicated by CGRP staining in cyan). Neurons activated by light brush stimulus are labelled with FOS staining (red). Scale bar: 100 μm. **h**. Quantification of Fluorogold+ cells co-labeled with FOS. n/N = # of sections/# of mice. **i**. Quantification of neuronal activation by SNI in spinal cord and supraspinal brain regions of the ascending nociceptive pathways. Activation level in sham-treated WT mice for each region is indicted by dashed lines. SC: spinal cord; LPBN: lateral parabrachial nucleus; mRt: mesencephalic reticular formation; Amy: Amygdala; MD: mediodorsal thalamus; VPL: central posterolateral thalamic nucleus; S2: secondary somatosensory cortex; S1HL: Primary somatosensory cortex, hindlimb region; ACC: anterior cingulate cortex; mPFC: medial prefrontal cortex. *, *p* < 0.05; **, *p* < 0.01; ***, *p* < 0.001. N = # of mice. **j**. Quantification of neuronal activation 1.5 hr after formalin injection in spinal cord and supraspinal brain regions of the ascending nociceptive pathways. *, *p* < 0.05; **, *p* < 0.01; ***, *p* < 0.001. N = # of mice. Statistical analysis was performed using two-sided unpaired t-test. Data shown as mean ± s.e.m.

Given that most SNI-induced FOS expression in spinal cord and brain regions occurs in non-PV-expressing neurons, we speculated that the RARα deletion in PV+ neurons blocks allodynic pain via changes at the circuit level. We first examined the laminal distribution of light-brush-activated FOS expression in the dorsal horn. We used calcitonin gene-related peptide (CGRP) staining to identify the lamina I and the outer lamina II (IIo) ^39-41^ (Fig. 2c). Most dorsal horn PV+ neurons are known to localize in lamina III^42^ (82.9%), while the inner lamina II (IIi) is the layer between the CGRP-positive lamina IIo and lamina III. Elevated light brush-induced FOS expression in SNI animals was observed in all laminae (I+IIo, IIi, III-VI) of the WT ipsilateral dorsal horn, but was significantly reduced across all laminae of the dorsal horn in PV-RARα cKOs (Fig. 2d).

Next, we used unilateral fluorogold microinjections into the lateral parabrachial nucleus (LPBN) to retrogradely label nociceptive projection neurons^40^, and quantified the extent of SNI-induced hyperactivation specifically in these neurons (Fig. 2e, f). Fluorogold labelling in the dorsal horn was concentrated in the superficial laminae (I+IIo), although a few fluorogold-labeled neurons were also found in lamina V^40^ (Fig. 2g). We observed comparable numbers of fluorogold-labelled neurons in lamina I+IIo of WT and PV-RARα cKO mice (WT: 83 vs cKO: 87, n = 3 mice/group). The majority of these projection neurons were activated by light-brush stimulation in WT mice (78.9% ± 4.3%) as assessed by FOS immunocytochemistry, but only a small fraction of these projection neurons were activated in PV-RARα cKO mice (27.6% ± 2.1%) (Fig. 2h). Thus, SNI caused an excessive activation of nociceptive projecting neurons by a light tactile stimulus, and the RARα deletion in PV+ neurons prevented this form of hyperactivation.

We asked whether similar changes in neuronal activation are observed in other regions of the central nociceptive pathway. Ipsilateral spinal nociceptive projection neurons cross to the contralateral spinal cord and ascend to the brain via the anterolateral tract^5^. In WT mice, SNI results in increased neuronal activation in response to light brush in both ipsilateral spinal and contralateral supraspinal regions of the ascending nociceptive pathway (Fig. 2i). Specifically, in addition to neurons in multiple laminae of the ipsilateral dorsal horn, neurons in the contralateral lateral parabrachial nucleus (LPBN), ventral posterolateral nucleus of thalamus (VPL), mesencephalic reticular nucleus (mRt), mediodorsal thalamus (MD), primary somatosensory cortex (S1 hind limb region), secondary somatosensory cortex (S2), anterior cingulate cortex (ACC), and medial prefrontal cortex (mPFC) all exhibited elevated FOS immunoreactivity (Fig. 2i), indicative of a general hyperactivation of nociceptive brain circuits. Ipsilateral S1, ACC and mPFC also showed significantly higher FOS expression compared to sham-lesion mice, reflecting the bilateral projections of pain and mechanosensory pathways in cortical regions (Fig. 2i). Strikingly, deletion of RARα in PV+ neurons prevented the hyperactivation of nociceptive pathway neurons throughout the CNS (Fig. 2i). Moreover, in most of the brain regions examined, the RARα deletion in PV+ neurons reduced their basal activity below the sham lesion level (dashed lines), indicating that the PV-RARα cKO not only prevented the development of hyperactivity in local circuits, but also dampened the basal tone of these circuits (Fig. 2i).

Finally, we examined responses to acute inflammatory pain induced by formalin injection into the hind paw. A comparable elevation of FOS expression was observed in the ipsilateral spinal dorsal horn and in contralateral brain regions in WT and PV-RARα cKO mice (Fig. 2j). Thus, RARα expression in PV+ neurons is specifically required for development of chronic neuropathic pain, but not for acute inflammatory pain.

### Dorsal horn PV+ neuron-specific deletion of RARα prevents mechanical allodynia

Our results up to this point show that global deletion of RARα in PV+ neurons prevents mechanical allodynia and reduces neuronal activity throughout central pain circuits. As one of four major inhibitory interneurons in the dorsal horn, PV+ neurons gate signals from low-threshold mechanoreceptive (LTMR) afferents to postsynaptic excitatory interneurons during tactile perception^17^. One possibility is that SNI modifies the local circuits involving PV+ neurons, thereby causing a misdirection of the tactile signal into the nociceptive pathway, and that deletion of RARα in dorsal horn PV+ neurons blocks this local circuit modification and prevents the induction of mechanical allodynia. Alternatively, deletion of RARα in PV+ neurons in cortical and subcortical brain regions may alter cortical outputs, which provide an essential descending feedback signal that blocks spinal cord hyperactivity^43,44^. To distinguish between these two possibilities, we crossed PV-Flp driver mice with floxed RARα cKO mice, and injected an adeno-associated virus encoding Flp-dependent Cre and tdTomato into the dorsal horn. In this manner, we selectively deleted RARα only in dorsal horn PV+ neurons. The Flp-targeting fidelity, quantified as percent of tdTomato+ neurons among PV+ neurons, was ∼84%, while the PV+ neuron targeting efficacy in the dorsal horn was ∼75% (Fig. 3a-c). Three weeks after viral infection, we performed SNI surgery, and tested mechanical allodynia starting three days after the surgery.

**Fig. 3.**
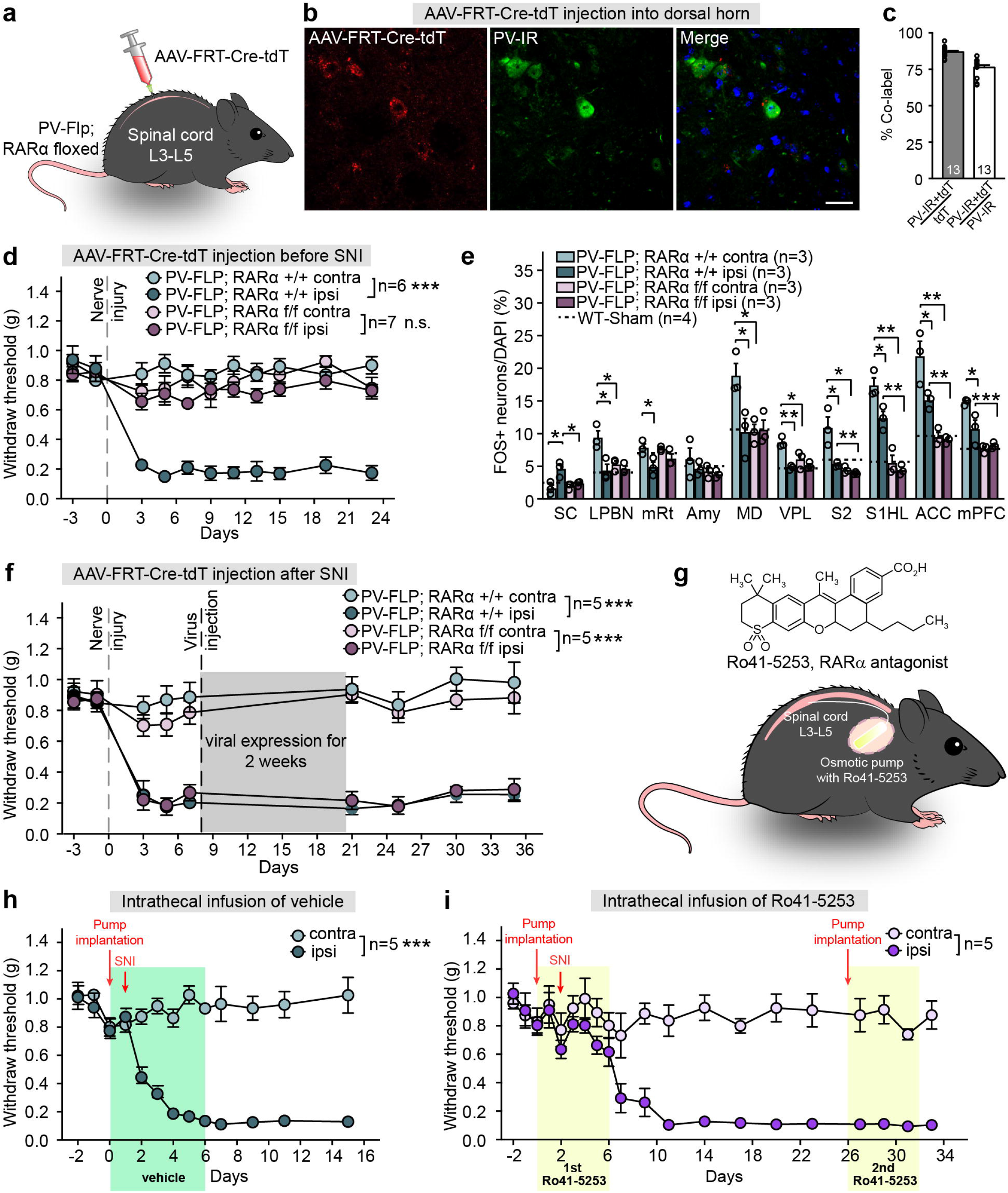
Region-specifical deletion of PV-RARα prevents the development of neuropathic pain. **a**. Experimental design to obtain RARα deletion in dorsal horn PV+ neurons. **b**. Representative images showing FRT-Cre-tdTomato expression in PV-immunoreactive neurons in spinal dorsal horn. Scale bar: 20 μm. **c**. Quantification of PV-Flp driver line expression fidelity (% PV-IR/tdTomato) and efficacy (% tdTomato/PV-IR) in spinal dorsal horn. **d**. Mechanical allodynia quantified as paw withdrawal threshold with von Frey filaments in WT and spinal-specific PV-RARα cKO mice before and over a period of three weeks after SNI. ***, *p* < 0.001. Two-way ANOVA with Bonferroni test. **e**. Quantification of neuronal activation by SNI in spinal cord and supraspinal brain regions of the ascending nociceptive pathways. Activation level in sham-treated WT mice for each region is indicted by dashed lines. SC: spinal cord; LPBN: lateral parabrachial nucleus; mRt: mesencephalic reticular formation; Amy: Amygdala; MD: mediodorsal thalamus; VPL: ventral posterolateral thalamic nuclei; S2: secondary somatosensory cortex; S1HL: Primary somatosensory cortex, hindlimb region; ACC: anterior cingulate cortex; mPFC: medial prefrontal cortex. *, *p* < 0.05; **, *p* < 0.01; ***, *p* < 0.001, two-sided unpaired t-test. N = # of mice. **f**. Mechanical allodynia quantified as paw withdrawal threshold with von Frey filaments in WT and spinal-specific PV-RARα cKO mice before and after SNI. AAV-FRT-Cre-tdT was injected a week after SNI. ***, *p* < 0.001, two-way ANOVA with Bonferroni test. **g**. Cartoon diagram showing implantation of osmotic pump with RARα antagonist Ro41-5253 and infusion site in the spinal cord. **h**. Mechanical allodynia quantified as paw withdrawal threshold in animals receiving vehicle infusion and SNI. ***, *p* < 0.001. Two-way ANOVA with Bonferroni test. **i**. Mechanical allodynia quantified as paw withdrawal threshold in animals receiving two infusions of a RARα antagonist Ro41-5253 (6-7 days each) and SNI. First infusion started 2 days before SNI and second infusion started 24 days after SNI when mechanical allodynia was evident. First infusion: *p* = 0.1; second infusion: ***, *p* < 0.001, two-way ANOVA followed by Bonferroni test. Data shown as mean ± s.e.m.

WT mice exhibited SNI-induced mechanical allodynia that lasted for the duration of observation (a minimum of 3 weeks). However, we observed no allodynia in mice with the spinal PV-RARα cKO, suggesting a critical involvement of RARα expression in dorsal horn PV+ neurons in the development of neuropathic pain (Fig. 3d). Additionally, SNI-induced neuronal hyperactivity in ascending nociceptive pathways (ipsilateral spinal dorsal horn, contralateral LPBN, mRt, MD, VPL, S2, S1HL, ACC and mPFC) was also prevented by the spinal PV-RARα cKO (Fig. 3e). Moreover, unlike the global PV-RARα deletion, the spinal cord-specific RARα deletion did not affect the basal activity of neurons in any of the regions examined as assayed by FOS expression (Fig. 3e). These results indicate that the reduced basal activity in brain regions in the global PV-RARα cKO is not the main cause of blocked hyperactivity in the brain regions involved in neuropathic pain. The specific involvement of spinal cord PV-RARα expression in allodynia was further verified by region-specific PV-RARα deletions in the ACC and S1, both of which were ineffective in preventing the development of SNI-induced mechanical allodynia (Extended Data Fig. S4).

Can the deletion of RARα in PV+ neurons reverse existing mechanical allodynia? To address this critical question, we injected the AAVs encoding Flp-dependent Cre and tdTomato into the dorsal horn of mice one week after SNI surgery, when mechanical allodynia is fully developed. Two weeks after vial infection, both WT and cKO mice showed robust mechanical allodynia for an additional two weeks (Fig. 3f), indicating that RARα is critically involved in the induction of circuit plasticity that establishes SNI-induced mechanical allodynia, but is not required for its maintenance.

### Spinal RARα antagonist infusion prevents mechanical allodynia

Given the essential function of RARα in dorsal horn PV+ neurons in the development of mechanical allodynia, we next asked whether other means of inhibiting RARα could prevent neuropathic pain development. Towards this end, we intrathecally delivered the selective RARα antagonist Ro41-5253 for seven days with an osmotic pump at the lumbar vertebrae L3-5 ^45,46^ (Fig. 3g). SNI surgery was performed two days after pump implantation. SNI-induced mechanical allodynia was detected in the vehicle infusion group for at least two weeks after surgery, indicating that the pump implantation surgery does not interfere with allodynia development (Fig. 3h). Infusion of Ro41-5253 for two days before the SNI surgery did not change tactile perception, as evidenced by the unchanged withdraw threshold in response to innocuous mechanical stimulation (Fig. 3i). However, continuous infusion of Ro41-5253 for an additional five days prevented the development of SNI-induced mechanical allodynia during this period (Fig. 3i, first infusion), indicating that locally blocking spinal RARα before nerve injury effectively prevents SNI-induced mechanical allodynia. To further test the potential impact of the RARα antagonist on existing allodynia, we allowed neuropathic pain to develop after the first pump was emptied for three more weeks, and infused Ro41-5253 again. We found that the RARα antagonist did not reverse the existing allodynia (Fig. 3i, second infusion). Taken together, these data confirm that blocking spinal RARα prior to, but not after, nerve injury prevents the development of mechanical allodynia, indicating that RARα functions within a critical time window after nerve injury to induce plastic changes in spinal circuits that lead to central sensitization and chronic neuropathic pain.

### RARα is required in PV+ neurons for SNI-induced disinhibition in spinal circuit

Our results so far strongly suggest a spinal cord mechanism involving PV+ neurons that drives development of SNI-induced mechanical allodynia. Two potential mechanisms have been proposed that involve changes in intrinsic membrane excitability or synaptic connectivity of PV+ neurons^9,17^.

We first investigated the contribution of RARα to SNI-induced changes in PV+ neuron excitability. We patched visually identified PV+ neurons in the dorsal horn, and verified their identity via their characteristic tonic fast-spiking firing patterns in response to current injections^9,17,42^ (Extended Data Fig. 5a, b). We observed no changes in passive membrane properties (resting membrane potential, membrane capacitance and input resistance) resulting from SNI or RARα deletion (Extended Data Fig. 5e-g). Consistent with previous findings^9^, SNI reduced the membrane excitability and increased the action-potential rheobase of PV+ neurons on the ipsilateral (lesion), but not the contralateral side (Extended Data Fig. 5c, d). Interestingly, we observed similar changes in membrane excitability in RARα cKO PV+ neurons (Extended Data Fig. 5c, d), suggesting that RARα is not required for SNI-induced changes in membrane properties. Consistent with these findings in dorsal horn PV+ neurons, ACC PV+ neurons that were identified visually and confirmed electrophysiologically (Extended Data Fig. 6a, b) also showed a reduced excitability and an increased AP rheobase on the contralateral side compared to the ipsilateral side (Extended Data Fig. 6c, d), without changes in passive membrane properties (Extended Data Fig. 6e-g). Again, deletion of PV-RARα did not block these SNI-induced changes in neuronal excitability (Extended Data Fig. 6c – g). Taken together, these results rule out membrane excitability changes as the main mechanism underlying RARα-dependent mechanical allodynia development.

We next investigated potential synaptic mechanisms for SNI-induced allodynia in the spinal cord. Two distinct forms of inhibition, presynaptic axoaxonic inhibition on LTMRs and postsynaptic inhibition of excitatory interneurons, have been described for dorsal horn PV+ neurons^9^. Dorsal horn PV+ neurons are thought to gate mechanical pain perception through both forms of synaptic inhibition that target multiple interneurons in lamina II, including SST+ cells,^47^ PKCγ+ cells^17,48,49^ and vGluT3+ cells^50,51^. Thus, an altered PV+ inhibitory synaptic output could plausibly underlie mechanical allodynia. To selectively activate PV+ synaptic outputs, we expressed channelrhodopsin (ChR2) in PV+ neurons by crossing a PV-Cre driver line with a floxed-ChR2 line. We then assayed presynaptic inhibition by utilizing a paradoxical property of this synapse, namely the GABA-mediated primary afferent depolarization (PAD) at lower temperature^9,52,53^ (Fig. 4a). Specifically, at room temperature (23°C) GABA release from PV+ neuron terminals depolarizes afferent terminals at axoaxonic synapses, thereby causing glutamate release that generates excitatory postsynaptic currents (EPSCs).

**Fig. 4.**
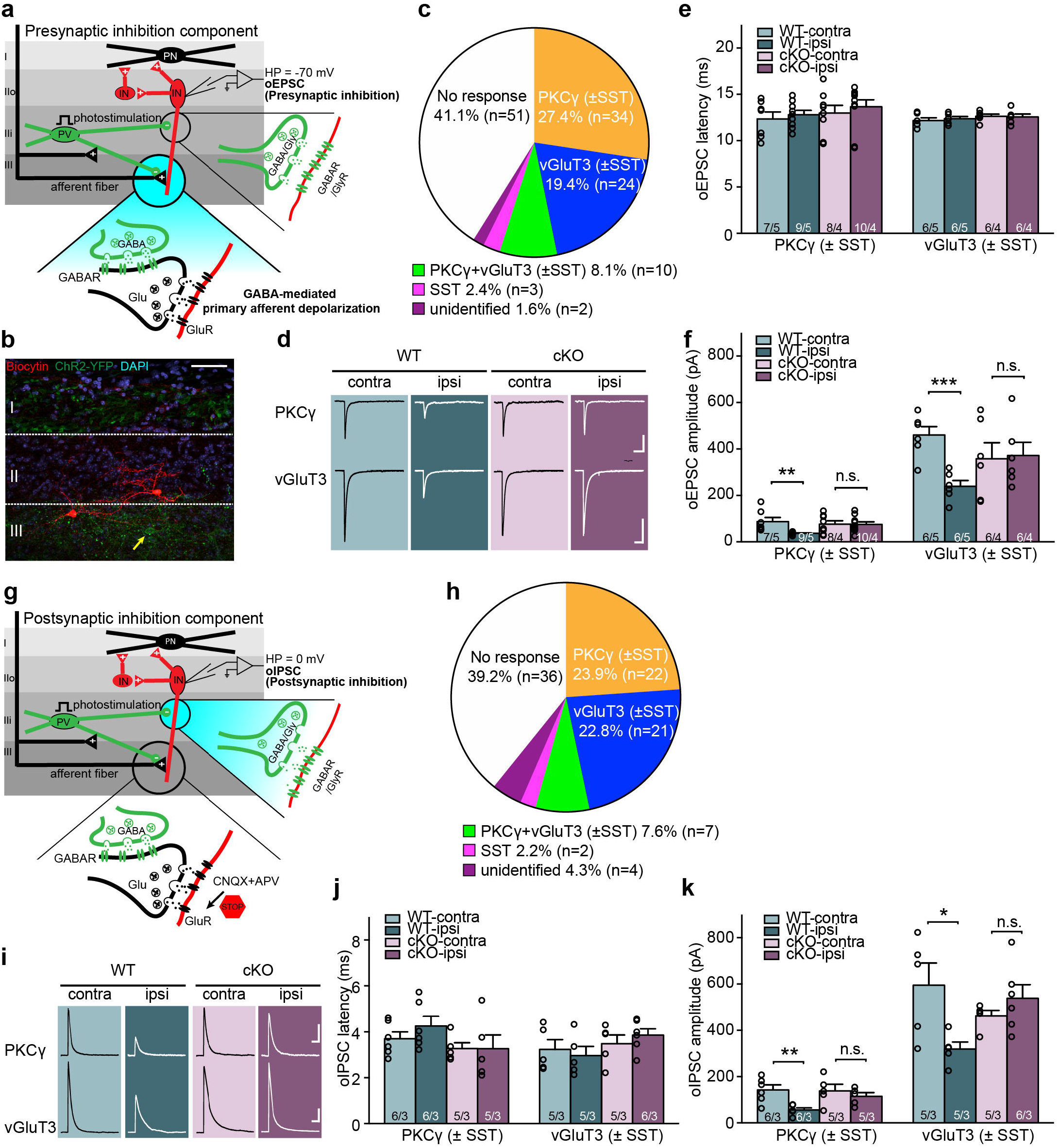
Deletion of RARα in PV+ cells prevents the SNI-induced presynaptic and postsynaptic disinhibition. **a**. Schematic diagram of local spinal dorsal horn circuit. PV+ neurons (green) gate primary afferent excitation onto excitatory postsynaptic interneurons (red) via synaptic inhibition onto both primary afferent terminals (indirect, presynaptic inhibition) and excitatory interneurons (direct, postsynaptic inhibition). Blue highlighted part depicts presynaptic inhibition component of the gate. Optogenetic activation of PV+ neurons induces oEPSC in postsynaptic interneurons via primary afferent depolarization (PAD), which triggers glutamate release from afferent terminals at room temperature. oEPSC is recorded with postsynaptic interneuron held at -70 mV (reversal potential of IPSC). **b**. Representative images shows that biocytin-labelled cells (red) in substantia gelatinosa. Yellow arrow indicates a PV-ChR2-YFP cell. Scale bar: 50 μm. **c**. Single cell qRT-PCR-based categorization of all neurons recorded for PV+ presynaptic inhibition measurement. **d**. Samples traces of oEPSCs recorded from PKCγ+ and vGluT3+ postsynaptic neurons in WT and cKO mice bilaterally. Scale bars: 100 ms, 50 pA (top), 200 pA (bottom). **e**. Synaptic latency of oEPSC in both WT and cKO mice. n/N = # of neurons/# of mice. **f**. Quantification of oEPSC amplitudes from PKCγ+ and vGluT3+ postsynaptic neurons in response to optogenetic activation of PV+ neurons. **, *p* < 0.01; ***, *p* < 0.001; two-sided unpaired t-test. n/N = # of neurons/# of mice. **g**. Schematic diagram of local spinal dorsal horn circuit with postsynaptic inhibitory gate highlighted in blue. oIPSCs are measured in postsynaptic interneurons (held at 0 mV) with optogenetic activation of PV+ cells in the presence of CNQX and APV. **h**. Single cell qRT-PCR-based categorization of all neurons recorded for PV+ postsynaptic inhibition measurement. **i**. Sample traces of oIPSCs recorded from PKCγ+ and vGluT3+ postsynaptic neurons in WT and cKO mice bilaterally. Scale bars: 100 ms, 50 pA (top), 100 pA (bottom). **j**. Synaptic latency of oIPSC in both WT and cKO mice. n/N = # of neurons/# of mice. **k**. Quantification of oIPSC amplitudes from PKCγ+ and vGluT3+ postsynaptic neurons in response to optogenetic activation of PV+ neurons. *, *p* < 0.05; **, *p* < 0.01, two-sided unpaired t-test. n/N = # of neurons/# of mice. Data shown as mean ± s.e.m.

A week after SNI surgery, we recorded from lamina IIneurons, held the neurons at the reversal potential of IPSCs, and recorded optogenetically evoked EPSCs (oEPSCs) in response to photo-stimulation of PV-ChR2 cells (Fig. 4b, Extended Data Fig. 7a, b). Full-field photostimulation of PV-ChR2-expressing neurons was achieved using single light pulses. To control for variabilities in light penetration and cell depth in slices, we used maximum synaptic responses generated by saturating light intensities for comparison (Extended Date Fig. 7c). Cells were selected from both ipsilateral and contralateral sides to achieve within-subject controls. Among 124 recorded neurons (56 in WT and 68 in RARα cKO, Fig. 4c), 19 neurons in WT (10 contralateral and 9 ipsilateral) and 32 neurons in RARα cKO (18 contralateral and 14 ipsilateral) did not show any evoked response, indicating a comparable level of PV-target connectivity in both WT and RARα cKO groups. The remaining 73 neurons (37 in WT and 36 in RARα cKO) showed clear synaptic responses that could be completely blocked by the AMPA-receptor antagonist CNQX (Fig. 4c and Extended Data Fig. 7d). To resolve the molecular identity of these recorded cells (n = 73), we recovered their RNA and performed single-cell qRT-PCR. These experiments revealed that among the postsynaptic target neurons of PV+ cells in the dorsal horn, PKCγ- and vGluT3-expressing cells are the two major types of excitatory interneurons and represented the majority of the PV-responsive neurons (∼ 80%) we recorded (Fig. 4c). The small number of remaining cells were either positive (10) or negative for both PKCγ and vGluT3 (5).

We analyzed the oEPSCs in the PKCγ+ and vGluT3+ neurons as they represent the majority of the recorded population. In general, the oEPSCs of both types of neurons showed similar long synaptic latencies (Fig. 4d, e), indicative of polysynaptic responses^9^. The response amplitudes were greater in vGluT3+ neurons than in PKCγ+ neurons (Fig. 4d, f), suggesting differences in target-specific connection strength. In WT neurons, ipsilateral oEPSC amplitudes in both PKCγ+ and vGluT3+ neurons were significantly reduced by SNI, indicating a weakened presynaptic inhibition by PV+ neurons compared to the contralateral control side (Fig. 4d, f). Importantly, this reduction was blocked by the PV-RARα cKO (Fig. 4d, f). Taken together, these data reveal that PV+ neuron-mediated presynaptic disinhibition induces, at least in part, the hypersensitivity of spinal pain circuits underlying mechanical allodynia, and that this SNI-induced synaptic change requires RARα expression in the PV+ neurons.

To assay the direct postsynaptic inhibition by PV+ neurons, we performed oIPSC recordings in acute spinal cord slices in the presence of CNQX and APV, which blocks all excitatory synaptic transmission (Fig. 4g and Extended Data Fig. 7e). We recorded from a total of 92 cells (42 WT and 50 cKO), among which 14 WT cells (7 control side, 7 lesion side) and 22 cKO cells (12 control side and 10 lesion side) did not show any responses, again exhibiting a similar degree of synaptic connectivity among all groups (Fig. 4h). In the remaining 56 responsive neurons, we also observed a majority of PKCγ+ (22) and vGluT3+ (21) neurons (Fig. 4h). Unlike presynaptic inhibition-induced oEPSCs, direct postsynaptic oIPSCs exhibited a shorter synaptic response latency, which was comparable across all groups (Fig. 4j). oIPSCs in vGluT3+ neurons were greater in amplitude compared to those in PKCγ+ neurons, and both types of neurons showed reduced oIPSCs after SNI on the ipsilateral (lesion) side (Fig. 4i, k). Deletion of RARα in PV+ neurons did not alter oIPSCs in control neurons, but completely prevented the SNI-induced reduction in postsynaptic inhibition (Fig. 4k), indicating that SNI-induced PV+ neuron-mediated postsynaptic disinhibition is also RARα-dependent.

Together, our synaptic response analysis provides unequivocal evidence for concomitant pre- and post-synaptic disinhibition of PV+ output onto specific postsynaptic targets in the spinal dorsal horn circuit underlying SNI-induced central sensitization. Importantly, these synaptic changes resemble homeostatic plasticity found in other inhibitory synapses in the brain and share the same molecular mechanism, in which RARα expression in PV+ neurons is a key component.

## DISCUSSION

Homeostatic synaptic plasticity has been studied extensively in the context of maintaining neural network stability through counteracting and preventing runaway Hebbian plasticity^54-56^, but its physiological role and pathological actions are incompletely understood. We now show that peripheral nerve injury, which leads to a chronic aberration of afferent input signals to spinal cord circuits, triggers homeostatic synaptic compensation in local spinal dorsal horn neurons, and thereby, as an unwanted by-product with vast pathophysiological consequences, causes the redirection of tactile sensory signals into spinal pain circuits, thus inducing allodynic pain.

In the spinal dorsal horn, nociceptive and tactile mechanosensory information are transmitted to separate circuits comprising projection neurons and local interneurons^57-59^. Under normal conditions, an innocuous touch activates mechanosensory inputs via myelinated Aδ and Aβ fibers as well as unmyelinated C fibers, thereby exciting mechanosensory projection neurons and generating a touch sensation^60,61^. Normally, the flow of sensory information from Aβ fibers to nociceptive circuits is blocked by local inhibitory interneurons that potently inhibit excitatory interneurons, such as somatostatin-expressing interneurons, and additionally suppress presynaptic sensory terminals, thus preventing activation of nociceptive projection neurons and ascending nociceptive pathway by tactile sensory inputs under normal conditions.^17,41,51,62-64^. In this study, we show that the loss of afferent inputs reduces synaptic inhibition onto both primary afferent axonal terminals (presynaptic inhibition) and postsynaptic excitatory relay interneurons (postsynaptic inhibition) by a homeostatic mechanism requiring RARα in PV-expressing inhibitory interneurons. We demonstrate that blocking homeostatic synaptic plasticity in spinal PV+ neurons by the RARα deletion prevents the reduction of synaptic inhibition by SNI and the development of mechanical allodynia. These results not only show that homeostatic synaptic plasticity is a key mechanism driving allodynic pain, but also suggest that the homeostatic plasticity pathway could be a therapeutic target for neuropathic pain.

Several spinal dorsal horn interneuron subtypes have been implicated in central sensitization and mechanical allodynia, including interneurons that express somatostatin^41^, dynorphin^41^, calretinin^51^, vesicular glutamate transporter vGluT351, and PV^17^. In the brain, PV-expressing GABAergic interneurons effectively control the spike patterns of pyramidal neurons by synapsing onto their axon initial segment (PV-expressing chandelier cells) or onto their soma and proximal dendrites (basket cells)^65-69^. In the spinal dorsal horn, PV+ neurons (75% of which express GABA and glycine^70^) primarily localize to the inner lamina II and lamina III^42^, where they provide an additional mode of synaptic inhibition termed presynaptic inhibition^24^. By forming axo-axonic synapses onto primary afferent terminals, PV+ neurons activate GABAARs present in the afferent terminals and curtail glutamate release from primary afferent inputs through shunting inhibition^42^. Spinal PV+ neurons normally tightly limit the flow of tactile sensory information from peripheral inputs to ascending nociceptive pathways via this form of presynaptic inhibition and via their postsynaptic inhibition of excitatory vGluT3+ or PKCγ+ interneurons ^62,71^. Here, we found that the inhibition of PKCγ+ and vGluT3+ interneurons by PV+ neurons is significantly reduced after peripheral nerve injury, illustrating the widespread impact of postsynaptic inhibition in the context of neuropathic pain. Furthermore, we demonstrate that SNI reduces presynaptic inhibition onto primary afferents, which synapse onto both PKCγ+ and vGluT3+ interneurons. PKCγ+ and vGluT3+ interneurons, the major excitatory interneuron subtypes in dorsal horn laminae II and III that relay Aβ afferent input to lamina I nociceptive projection neurons, have both been implicated in neuropathic pain ^17,48,50,72,73^. The two concomitantly occurring effects of inhibitory synaptic plasticity onto PKCγ+ and vGluT3+ neurons described in our study demonstrate that the spinal dorsal horn circuit adjusted itself homeostatically to compensate for the loss of peripheral input due to the nerve injury during SNI, leading to hypersensitivity of the ascending nociceptive pathway and neuropathic pain.

Compared to the circuit level remodeling in spinal dorsal horn, much less is understood about the molecular players enabling synaptic and circuit plasticity underlying neuropathic pain. In dissecting the molecular mechanisms underlying the homeostatic plasticity that induces disinhibition of sensory gating by PV+ neurons, we examined the role of RARα because of its central contribution to most forms of homeostatic synaptic plasticity in in the brain^30,31,33-35^. In mature neurons, RARα translocates from the nucleus to the cytosol to act as a retinoic acid-dependent translational regulator in homeostatic synaptic plasticity^74,75^. Synaptic RA/RARα signaling is essential for homeostatic synaptic plasticity at both excitatory and inhibitory synapses in the hippocampus and neocortex^30-34^. Of particular interest is RARα’s involvement in homeostatic synaptic plasticity in the primary visual cortex (V1). Prolonged visual deprivation induces homeostatic synaptic plasticity at both V1 excitatory and inhibitory cortical synapses toward increased synaptic excitation/inhibition balance ^36,76,77^. Among the synapses involved, synaptic inhibition from PV+ neurons to L2/3 pyramidal neurons shows profound reduction after loss of visual input, a form of homeostatic plasticity that requires RARα expression in PV+ neurons. In the current study, we demonstrate that a similar form of homeostatic synaptic plasticity occurs in spinal dorsal horn PV+ neurons, with RARα as the key effector. Peripheral nerve injury drives homeostatic changes that lead to a loss of inhibitory output from PV+ neurons, which causes primary afferent inputs to erroneously drive superficial lamina pain projecting neuron and ascending nociceptive pathway activation. Using PV-specific genetic deletion of RARα, this form of homeostatic synaptic plasticity is inhibited and the gate remains closed, thus preventing the development of mechanical allodynia. These results indicate that the basic molecular mechanisms underlying homeostatic synaptic plasticity may be conserved across CNS circuits. Pathological neuronal activity resulting from homeostatic plasticity after nerve injury or cochlear trauma is a common theme in many types of chronic pain or in tinnitus^78-80^. Thus, identifying key molecular players as potential druggable targets will be the first step towards identifying effective remedies for treating these debilitating conditions.

## METHODS SUMMARY

Full methods including animal breeding, SNI surgery, optogenetics and electrophysiology, in-situ hybridization, single-cell qRT-PCR, behavioral assessments, histology and statistics with associated references are available online.

## AUTHOR CONTRIBUTIONS

B.C. and L.C. designed the experiments and analyses. B.C. performed all the experiments and analyses with input from G.S. and L.C.. B.C. and L.C. wrote the manuscript with input from G.S..

## ACKNOWLEDGMENTS

We thank Drs. Dong Wang, Dan Berg and Chelsie L. Brewer for technical assistances, and Drs. Kristin Arendt and Omid Miry for helpful discussion of the study and critical reading of the manuscript. The work was supported by NIH grants MH086403 (L.C.), NS11566001 (L.C.), HD104458 (L.C.), and a Stanford School of Medicine Dean’s postdoctoral fellowship (B.C.).

## COMPETING INTERESTS

The authors declare no competing interests.

## SUPPLEMENTAL INFORMATION

Supplemental Information includes seven figures, which can be found with this article online at

<URL>.

## Notes

### Competing Interest Statement

The authors have declared no competing interest.

## Reference

1 Bouhassira, D. et al. Comparison of pain syndromes associated with nervous or somatic lesions and development of a new neuropathic pain diagnostic questionnaire (DN4). Pain 114, 29–36, doi:10.1016/j.pain.2004.12.010 (2005).

2 Mercer Lindsay, N., Chen, C., Gilam, G., Mackey, S. & Scherrer, G. Brain circuits for pain and its treatment. Science translational medicine 13, eabj7360, doi:10.1126/scitranslmed.abj7360 (2021).

3 von Hehn, C. A., Baron, R. & Woolf, C. J. Deconstructing the neuropathic pain phenotype to reveal neural mechanisms. Neuron 73, 638–652, doi:10.1016/j.neuron.2012.02.008 (2012).

4 Moehring, F., Halder, P., Seal, R. P. & Stucky, C. L. Uncovering the Cells and Circuits of Touch in Normal and Pathological Settings. Neuron 100, 349–360, doi:10.1016/j.neuron.2018.10.019 (2018).

5 Todd, A. J. Identifying functional populations among the interneurons in laminae I-III of the spinal dorsal horn. Molecular pain 13, 1744806917693003, doi:10.1177/1744806917693003 (2017).

6 Keller, A. F., Beggs, S., Salter, M. W. & De Koninck, Y. Transformation of the output of spinal lamina I neurons after nerve injury and microglia stimulation underlying neuropathic pain. Molecular pain 3, 27, doi:10.1186/1744-8069-3-27 (2007).

7 Torsney, C. & MacDermott, A. B. Disinhibition opens the gate to pathological pain signaling in superficial neurokinin 1 receptor-expressing neurons in rat spinal cord. J Neurosci 26, 1833–1843, doi:10.1523/JNEUROSCI.4584-05.2006 (2006).

8 Latremoliere, A. & Woolf, C. J. Central sensitization: a generator of pain hypersensitivity by central neural plasticity. The journal of pain 10, 895–926, doi:10.1016/j.jpain.2009.06.012 (2009).

9 Boyle, K. A. et al. Defining a Spinal Microcircuit that Gates Myelinated Afferent Input: Implications for Tactile Allodynia. Cell Rep 28, 526-+, doi:10.1016/j.celrep.2019.06.040 (2019).

10 Woolf, C. J., Shortland, P. & Coggeshall, R. E. Peripheral nerve injury triggers central sprouting of myelinated afferents. Nature 355, 75–78, doi:10.1038/355075a0 (1992).

11 Tulleuda, A. et al. TRESK channel contribution to nociceptive sensory neurons excitability: modulation by nerve injury. Molecular pain 7, 30, doi:10.1186/1744-8069-7-30 (2011).

12 Scholz, J. et al. Blocking caspase activity prevents transsynaptic neuronal apoptosis and the loss of inhibition in lamina II of the dorsal horn after peripheral nerve injury. The Journal of neuroscience : the official journal of the Society for Neuroscience 25, 7317–7323, doi:10.1523/JNEUROSCI.1526-05.2005 (2005).

13 Polgar, E., Gray, S., Riddell, J. S. & Todd, A. J. Lack of evidence for significant neuronal loss in laminae I-III of the spinal dorsal horn of the rat in the chronic constriction injury model. Pain 111, 144–150, doi:10.1016/j.pain.2004.06.011 (2004).

14 Ikeda, H., Heinke, B., Ruscheweyh, R. & Sandkuhler, J. Synaptic plasticity in spinal lamina I projection neurons that mediate hyperalgesia. Science 299, 1237–1240, doi:10.1126/science.1080659 (2003).

15 Gradwell, M. A., Callister, R. J. & Graham, B. A. Reviewing the case for compromised spinal inhibition in neuropathic pain. Journal of Neural Transmission 127, 481–503, doi:10.1007/s00702-019-02090-0 (2020).

16 Finnerup, N. B., Kuner, R. & Jensen, T. S. Neuropathic Pain: From Mechanisms to Treatment. Physiol Rev 101, 259–301, doi:10.1152/physrev.00045.2019 (2021).

17 Petitjean, H. et al. Dorsal Horn Parvalbumin Neurons Are Gate-Keepers of Touch-Evoked Pain after Nerve Injury. Cell Rep 13, 1246–1257, doi:10.1016/j.celrep.2015.09.080 (2015).

18 Peirs, C., Dallel, R. & Todd, A. J. Recent advances in our understanding of the organization of dorsal horn neuron populations and their contribution to cutaneous mechanical allodynia. Journal of neural transmission 127, 505–525, doi:10.1007/s00702-020-02159-1 (2020).

19 Lever, I., Cunningham, J., Grist, J., Yip, P. K. & Malcangio, M. Release of BDNF and GABA in the dorsal horn of neuropathic rats. Eur J Neurosci 18, 1169–1174, doi:10.1046/j.1460-9568.2003.02848.x (2003).

20 Fukuoka, T. et al. Change in mRNAs for neuropeptides and the GABA(A) receptor in dorsal root ganglion neurons in a rat experimental neuropathic pain model. Pain 78, 13–26, doi:10.1016/S0304-3959(98)00111-0 (1998).

21 Castro-Lopes, J. M., Malcangio, M., Pan, B. H. & Bowery, N. G. Complex changes of GABAA and GABAB receptor binding in the spinal cord dorsal horn following peripheral inflammation or neurectomy. Brain Res 679, 289–297, doi:10.1016/0006-8993(95)00262-o (1995).

22 Lu, Y. et al. A feed-forward spinal cord glycinergic neural circuit gates mechanical allodynia. J Clin Invest 123, 4050–4062, doi:10.1172/JCI70026 (2013).

23 Ibuki, T., Hama, A. T., Wang, X. T., Pappas, G. D. & Sagen, J. Loss of GABA-immunoreactivity in the spinal dorsal horn of rats with peripheral nerve injury and promotion of recovery by adrenal medullary grafts. Neuroscience 76, 845–858, doi:10.1016/s0306-4522(96)00341-7 (1997).

24 Comitato, A. & Bardoni, R. Presynaptic Inhibition of Pain and Touch in the Spinal Cord: From Receptors to Circuits. Int J Mol Sci 22, doi:10.3390/ijms22010414 (2021).

25 Yamamoto, T. & Yaksh, T. L. Effects of intrathecal strychnine and bicuculline on nerve compression-induced thermal hyperalgesia and selective antagonism by MK-801. Pain 54, 79–84, doi:10.1016/0304-3959(93)90102-U (1993).

26 Yaksh, T. L. Behavioral and autonomic correlates of the tactile evoked allodynia produced by spinal glycine inhibition: effects of modulatory receptor systems and excitatory amino acid antagonists. Pain 37, 111–123, doi:10.1016/0304-3959(89)90160-7 (1989).

27 Neumann, E., Kupfer, L. & Zeilhofer, H. U. The alpha2/alpha3GABAA receptor modulator TPA023B alleviates not only the sensory but also the tonic affective component of chronic pain in mice. Pain 162, 421–431, doi:10.1097/j.pain.0000000000002030 (2021).

28 Eaton, M. J., Martinez, M. A. & Karmally, S. A single intrathecal injection of GABA permanently reverses neuropathic pain after nerve injury. Brain Res 835, 334–339, doi:10.1016/s0006-8993(99)01564-4 (1999).

29 Juarez-Salinas, D. L. et al. GABAergic cell transplants in the anterior cingulate cortex reduce neuropathic pain aversiveness. Brain : a journal of neurology 142, 2655–2669, doi:10.1093/brain/awz203 (2019).

30 Park, E., Tjia, M., Zuo, Y. & Chen, L. Postnatal Ablation of Synaptic Retinoic Acid Signaling Impairs Cortical Information Processing and Sensory Discrimination in Mice. J Neurosci 38, 5277–5288, doi:10.1523/JNEUROSCI.3028-17.2018 (2018).

31 Zhong, L. R., Chen, X., Park, E., Sudhof, T. C. & Chen, L. Retinoic Acid Receptor RARalpha-Dependent Synaptic Signaling Mediates Homeostatic Synaptic Plasticity at the Inhibitory Synapses of Mouse Visual Cortex. J Neurosci 38, 10454–10466, doi:10.1523/JNEUROSCI.1133-18.2018 (2018).

32 Aoto, J., Nam, C. I., Poon, M. M., Ting, P. & Chen, L. Synaptic signaling by all-trans retinoic acid in homeostatic synaptic plasticity. Neuron 60, 308–320 (2008).

33 Sarti, F., Zhang, Z., Schroeder, J. & Chen, L. Rapid suppression of inhibitory synaptic transmission by retinoic acid. J Neurosci 33, 11440–11450 (2013).

34 Yee, A. X. & Chen, L. Differential regulation of spontaneous and evoked inhibitory synaptic transmission in somatosensory cortex by retinoic acid. Synapse 70, 445–452, doi:10.1002/syn.21921 (2016).

35 Hsu, Y. T., Li, J., Wu, D., Sudhof, T. C. & Chen, L. Synaptic retinoic acid receptor signaling mediates mTOR-dependent metaplasticity that controls hippocampal learning. Proc Natl Acad Sci U S A 116, 7113–7122, doi:10.1073/pnas.1820690116 (2019).

36 Zhong, L. R., Chen, X., Park, E., Sudhof, T. C. & Chen, X. Retinoic Acid Receptor RAR alpha-Dependent Synaptic Signaling Mediates Homeostatic Synaptic Plasticity at the Inhibitory Synapses of Mouse Visual Cortex. Journal of Neuroscience 38, 10454–10466, doi:10.1523/Jneurosci.1133-18.2018 (2018).

37 Decosterd, I. & Woolf, C. J. Spared nerve injury: an animal model of persistent peripheral neuropathic pain. Pain 87, 149–158, doi:Doi 10.1016/S0304-3959(00)00276-1 (2000).

38 Shields, S. D., Eckert, W. A. & Basbaum, A. I. Spared nerve injury model of neuropathic pain in the mouse: A behavioral and anatomic analysis. J Pain 4, 465–470, doi:10.1067/S1526-5900(03)00781-8 (2003).

39 Loken, L. S. et al. Contribution of dorsal horn CGRP-expressing interneurons to mechanical sensitivity. Elife 10, doi:10.7554/eLife.59751 (2021).

40 Wang, D. et al. Functional Divergence of Delta and Mu Opioid Receptor Organization in CNS Pain Circuits. Neuron 98, 90-+, doi:10.1016/j.neuron.2018.03.002 (2018).

41 Duan, B. et al. Identification of Spinal Circuits Transmitting and Gating Mechanical Pain. Cell 159, 1417–1432, doi:10.1016/j.cell.2014.11.003 (2014).

42 Hughes, D. I. et al. Morphological, neurochemical and electrophysiological features of parvalbumin-expressing cells: a likely source of axo-axonic inputs in the mouse spinal dorsal horn. Journal of Physiology-London 590, 3927–3951, doi:10.1113/jphysiol.2012.235655 (2012).

43 Liu, Y. et al. Touch and tactile neuropathic pain sensitivity are set by corticospinal projections. Nature 561, 547–550, doi:10.1038/s41586-018-0515-2 (2018).

44 Ossipov, M. H., Morimura, K. & Porreca, F. Descending pain modulation and chronification of pain. Current Opinion in Supportive and Palliative Care 8, 143–151, doi:10.1097/Spc.0000000000000055 (2014).

45 Ke, Q., Li, R., Cai, L., Wu, S. D. & Li, C. M. Ro41-5253, a selective antagonist of retinoic acid receptor alpha, ameliorates chronic unpredictable mild stress-induced depressive-like behaviors in rats: Involvement of regulating HPA axis and improving hippocampal neuronal deficits. Brain Research Bulletin 146, 302–309, doi:10.1016/j.brainresbull.2019.01.022 (2019).

46 Hu, P., van Dam, A. M., Wang, Y., Lucassen, P. J. & Zhou, J. N. Retinoic acid and depressive disorders: Evidence and possible neurobiological mechanisms. Neuroscience and Biobehavioral Reviews 112, 376–391, doi:10.1016/j.neubiorev.2020.02.013 (2020).

47 Zhang, Y. et al. Timing Mechanisms Underlying Gate Control by Feedforward Inhibition. Neuron 99, 941–955 e944, doi:10.1016/j.neuron.2018.07.026 (2018).

48 Malmberg, A. B., Chen, C., Tonegawa, S. & Basbaum, A. I. Preserved acute pain and reduced neuropathic pain in mice lacking PKCgamma. Science 278, 279–283, doi:10.1126/science.278.5336.279 (1997).

49 Neumann, S., Braz, J. M., Skinner, K., Llewellyn-Smith, I. J. & Basbaum, A. I. Innocuous, not noxious, input activates PKCgamma interneurons of the spinal dorsal horn via myelinated afferent fibers. The Journal of neuroscience : the official journal of the Society for Neuroscience 28, 7936–7944, doi:10.1523/JNEUROSCI.1259-08.2008 (2008).

50 Peirs, C. et al. Mechanical Allodynia Circuitry in the Dorsal Horn Is Defined by the Nature of the Injury. Neuron 109, doi:10.1016/j.neuron.2020.10.027 (2021).

51 Peirs, C. et al. Dorsal Horn Circuits for Persistent Mechanical Pain. Neuron 87, 797–812, doi:10.1016/j.neuron.2015.07.029 (2015).

52 Fink, A. J. et al. Presynaptic inhibition of spinal sensory feedback ensures smooth movement. Nature 509, 43–48, doi:10.1038/nature13276 (2014).

53 Rudomin, P. In search of lost presynaptic inhibition. Exp Brain Res 196, 139–151, doi:10.1007/s00221-009-1758-9 (2009).

54 Turrigiano, G. Homeostatic synaptic plasticity: local and global mechanisms for stabilizing neuronal function. Cold Spring Harbor perspectives in biology 4, a005736, doi:10.1101/cshperspect.a005736 (2012).

55 Monday, H. R., Younts, T. J. & Castillo, P. E. Long-Term Plasticity of Neurotransmitter Release: Emerging Mechanisms and Contributions to Brain Function and Disease. Annual Review of Neuroscience, Vol 41 41, 299–322, doi:10.1146/annurev-neuro-080317-062155 (2018).

56 Yee, A. X., Hsu, Y. T. & Chen, L. A metaplasticity view of the interaction between homeostatic and Hebbian plasticity. Philosophical transactions of the Royal Society of London. Series B, Biological sciences 372, doi:10.1098/rstb.2016.0155 (2017).

57 Todd, A. J. Neuronal circuitry for pain processing in the dorsal horn. Nat Rev Neurosci 11, 823–836, doi:10.1038/nrn2947 (2010).

58 Choi, S. et al. Parallel ascending spinal pathways for affective touch and pain. Nature 587, 258–263, doi:10.1038/s41586-020-2860-1 (2020).

59 Peirs, C. & Seal, R. P. Neural circuits for pain: Recent advances and current views. Science 354, 578–584, doi:10.1126/science.aaf8933 (2016).

60 Koch, S. C., Acton, D. & Goulding, M. Spinal Circuits for Touch, Pain, and Itch. Annu Rev Physiol 80, 189–217, doi:10.1146/annurev-physiol-022516-034303 (2018).

61 Handler, A. & Ginty, D. D. The mechanosensory neurons of touch and their mechanisms of activation. Nat Rev Neurosci 22, 521–537, doi:10.1038/s41583-021-00489-x (2021).

62 Hughes, D. I. & Todd, A. J. Central Nervous System Targets: Inhibitory Interneurons in the Spinal Cord. Neurotherapeutics 17, 874–885, doi:10.1007/s13311-020-00936-0 (2020).

63 Zholudeva, L. V. et al. Spinal Interneurons as Gatekeepers to Neuroplasticity after Injury or Disease. J Neurosci 41, 845–854, doi:10.1523/Jneurosci.1654-20.2020 (2021).

64 Takazawa, T. & MacDermott, A. B. Synaptic pathways and inhibitory gates in the spinal cord dorsal horn. Ann N Y Acad Sci 1198, 153–158, doi:10.1111/j.1749-6632.2010.05501.x (2010).

65 Cobb, S. R. et al. Synaptic effects of identified interneurons innervating both interneurons and pyramidal cells in the rat hippocampus. Neuroscience 79, 629–648, doi:10.1016/s0306-4522(97)00055-9 (1997).

66 Pawelzik, H., Hughes, D. I. & Thomson, A. M. Physiological and morphological diversity of immunocytochemically defined parvalbumin- and cholecystokinin-positive interneurones in CA1 of the adult rat hippocampus. The Journal of comparative neurology 443, 346–367, doi:10.1002/cne.10118 (2002).

67 Cobb, S. R., Buhl, E. H., Halasy, K., Paulsen, O. & Somogyi, P. Synchronization of neuronal activity in hippocampus by individual GABAergic interneurons. Nature 378, 75–78, doi:10.1038/378075a0 (1995).

68 Miles, R., Toth, K., Gulyas, A. I., Hajos, N. & Freund, T. F. Differences between somatic and dendritic inhibition in the hippocampus. Neuron 16, 815–823, doi:10.1016/s0896-6273(00)80101-4 (1996).

69 Lewis, D. A. & Lund, J. S. Heterogeneity of chandelier neurons in monkey neocortex: corticotropin-releasing factor- and parvalbumin-immunoreactive populations. The Journal of comparative neurology 293, 599–615, doi:10.1002/cne.902930406 (1990).

70 Abraira, V. E. et al. The Cellular and Synaptic Architecture of the Mechanosensory Dorsal Horn. Cell 168, 295–310 e219, doi:10.1016/j.cell.2016.12.010 (2017).

71 Braz, J., Solorzano, C., Wang, X. & Basbaum, A. I. Transmitting pain and itch messages: a contemporary view of the spinal cord circuits that generate gate control. Neuron 82, 522–536, doi:10.1016/j.neuron.2014.01.018 (2014).

72 Seal, R. P. et al. Injury-induced mechanical hypersensitivity requires C-low threshold mechanoreceptors. Nature 462, 651–655, doi:10.1038/nature08505 (2009).

73 Cheng, L. et al. Identification of spinal circuits involved in touch-evoked dynamic mechanical pain. Nat Neurosci 20, 804–814, doi:10.1038/nn.4549 (2017).

74 Maghsoodi, B. et al. Retinoic acid regulates RARalpha-mediated control of translation in dendritic RNA granules during homeostatic synaptic plasticity. Proc Natl Acad Sci U S A 105, 16015–16020 (2008).

75 Poon, M. M. & Chen, L. Retinoic acid-gated sequence-specific translational control by RARalpha. Proc Natl Acad Sci U S A 105, 20303–20308 (2008).

76 Miska, N. J., Richter, L. M., Cary, B. A., Gjorgjieva, J. & Turrigiano, G. G. Sensory experience inversely regulates feedforward and feedback excitation-inhibition ratio in rodent visual cortex. Elife 7, doi:10.7554/eLife.38846 (2018).

77 Maffei, A. & Turrigiano, G. G. Multiple modes of network homeostasis in visual cortical layer 2/3. The Journal of neuroscience : the official journal of the Society for Neuroscience 28, 4377–4384, doi:10.1523/JNEUROSCI.5298-07.2008 (2008).

78 Flor, H. et al. Phantom-limb pain as a perceptual correlate of cortical reorganization following arm amputation. Nature 375, 482–484, doi:10.1038/375482a0 (1995).

79 Eggermont, J. J. & Roberts, L. E. The neuroscience of tinnitus. Trends Neurosci 27, 676–682, doi:10.1016/j.tins.2004.08.010 (2004).

80 Moller, A. R. Tinnitus and pain. Prog Brain Res 166, 47–53, doi:10.1016/S0079-6123(07)66004-X (2007).

