## Supplementary methods and figures for "Spinal cord homeostatic plasticity gates mechanical allodynia in chronic pain"

### Materials and Methods

#### Animals

All animals were housed and used in experiments following Stanford University APLAC guidelines. Postnatal day 35-56 male and female littermates were used for this study. Mice were group-housed with littermates and maintained under a 12 h light/dark cycle. The RARα$^{fl/fl}$ mice (C57BL/6 background) were originally obtained as a generous gift from Dr. Pierre Chambon and Norbert Ghyselinck (IGBNC, Stasbourg, France)[1]. C57BL/6 wild-type mice were obtained from the Jackson Laboratory.

Cell type-specific RARα cKO mice were generated by crossing the RARα$^{fl/fl}$ line with cell type-specific driver lines (PV-Cre, stock 008069; SST-Cre, stock 013044, Jackson Laboratory, Bar Harbor, ME) and a EYFP reporter line (Stock 007903, Jackson Laboratory, Bar Harbor, ME). Litters were genotyped by PCR using the protocol described previously[2,3].

Primers used for Cre are:

Forward 5’-CACCCTGTTACGTATAGCCG-3’,

Reverse 5’-GAGTCATCCTTAGCGCCGTA-3’;

Primers for flox site of RARα are:

Forward 5’-GTGTGTGTGTGTATTCGCGTGC-3’,

Reverse 5’-ACAAAGCAAGGCTTGTAGATGC-3’;

Primers used for EYFP are:

WT-Forward: 5’-AAGGGAGCTGCAGTGGAGTA-3’,

WT-Reverse: 5'-CCGAAAATCTGTGGGAAGTC-3',
Mutant (Mu)-Forward: 5'-ACATGGTCCTGCTGGAGTTC-3',
Mu-Reverse: 5'-GGCATTAAAGCAGCGTATCC-3';
Primers for channelrhodopsin-2:
Mutant Forward: 5'-CAT TGG TGG CAC TGA GAT TG-3'
Mutant Reverse: 5'-GAA CTT CAG GGT CAG CTT GC-3'
Wild type Forward: 5'-AAG GGA GCT GCA GTG GAG TA-3'
Wild type Reverse: 5'-CCG AAA ATC TGT GGG AAG TC-3'

For regional ablation of RARα, RARα$^{fl/fl}$ mice were crossed to PV-2A-FlpO line (stock 022730, Jackson Laboratory, Bar Harbor, ME). WT mice crossed to PV-2A-FlpO served as controls. Litters were genotyped for flox and FlpO.
Primers used for FlpO are:
WT forward: 5'-GGA TGC TTG CCG AAG ATA AG-3',
Mu forward: 5'-CTG AGC AGC TAC ATC AAC AGG-3',
common reverse: 5'-TGT TTC TCC AGC ATT TCC AG-3'.

**In situ hybridization**

*In situ* hybridization with RNAscope Technology[4] (Advanced Cell Diagnostics Bioscience) was used to detect the expression of RARα mRNA. P35-P42 WT and RARα KO mice were deeply anesthetized with ketamine-xylazine followed by perfusion with 0.1M PBS and fixed with 4% PFA. Spinal cord L3-L5 regions were dissected, cryoprotected in 30% sucrose overnight, sectioned at 14 μm and maintained at -80 °C. On the day of ISH, tissue was thawed from -80 °C, washed with PBS and processed according to the manufacturer's protocol. Briefly, the tissue was permeabilized, incubated with protease for 30 min, and hybridized with probe(s) for 2 hr at 40 °C. After amplification of mRNA signals, the slides were mounted with DAPI. A scanning microscope (BX61VS; OLYMPUS) and a confocal microscope (Nikon Instruments) were used to obtain whole-spinal cord slice images and higher-resolution images, respectively.

**Sigle-cell qRT-PCR**

mRNAs extracted from single cells were amplified using the protocol previously described[5]. Acute slices were obtained from P35-P42 mice and single cell contents were extracted from the substantia gelatinosa in the spinal cord. mRNAs from single cell extracts were amplified use Superscript III One-Step RT-PCR System with Platinum Taq High Fidelity DNA polymerase (Invitrogen, cat 12574035). RT-qPCR was then performed using TaqMan Gene Expression Master Mix (Applied Biosystems, cat 4369016) with Taqman primers from Life Technologies; RARα (Mm00436262_m1), Actinβ (Mm02619580_g1), GFAP (Mm01253033_m1), PV (Mm00443100_m1), SST (Mm.PT.53a.7678291), PKCγ (Mm.PT.58.45983184), VGluT3 (Mm.PT.58.17676720). Actinβ was used as the endogenous control. Cells expressing GFAP, a marker for astrocytes, were excluded from the sample.

**Spared nerve injury surgery**

The spared nerve injury (SNI) surgery was performed based on previously established methods[6]. Briefly, mice were anesthetized with a mixture of ketamine and xylazine (0.1 ml/20g, intraperitoneal injection). The sciatic nerve with three branches was exposed, and the common peroneal and tibial branches were ligated with 8-0 nylon suture, sectioned distal to the ligation, removing 2-4 mm of the distal nerve stump. Care was taken to avoid any stretching or contact with the spared sural nerve. Sham operation without axotomy was served as control.

**Osmotic pump implantation**

P35-P42 mice were anesthetized with 2.5% isoflurane, placed into a stereotaxic frame and a polyethylene catheter (Mouse Intrathecal Catheter MIT-02) was inserted into the spinal subarachnoid space of each mouse so as to place its tip 1-1.5 mm rostral to the dural incision at the L3-5 intervertebral space. The other end of the catheter was attached to an osmotic pump (Alzet 1007D) that infused the solution at a flow rate of 0.5 µl/h for 7 days. The osmotic pump was filled with Ro41-5253 (1.25 µg/h) or vehicle (aCSF). All animals had free access to food and water from the time of surgery until they were sacrificed. Each mouse was examined every day for behavioral changes.

**AAV Vector construction, AAV preparation and Stereotaxic injections**

The AAV-Frt-Cre-tdTomato vector was constructed using a synapsin promoter that drives the expression of an inverted Cre-IRES-tdTomato sequence, which is flanked by two FRT F545 sites. The AAV Frt-Cre-tdTomato was packaged with AAV-DJ capsids for high efficiency in vivo neuronal infection. AAVs were prepared as described[7]. For intraspinal injections, a midline incision along the left lumbar vertebrae was carefully performed until the spinal cord was visible from the intervertebral spaces. No laminectomy was performed to maximally avoid trauma. Two stereotaxic injections of 250 nl AAVs on each side 400 μM lateral to the posterior spinal arteries and 400 μm from the dura were made in L3-5. The lassimus dorsi were sutured to protect the spinal cord and the skin was sealed with silk sutures[8]. For intracortical injections, 400 nL AAVs were injected into the anterior cingulate cortex (in mm: AP 0.90/1.40, ML ± 0.25, DV 1.50) or somatosensory cortex hindlimb region (in mm: AP -0.30/-0.80, ML ±1.50, DV 1.00)[9,10].

**Retrograde labeling**

7 days after SNI surgery, WT or PV-Cre RARα KO mice were anaesthetized with isofluorane and 300 nl of fluorogold (FG, 2% in sterilized water) were injected unilaterally with a glass micropipette into the lateral parabrachial nucleus (AP 5.0 mm; ML 1.4 mm; DV -2.3 mm from brain surface). 5-7 days following injection, spinal cord and brain tissues was collected for histological analysis.

**Immunohistochemistry**

**Innocuous mechanical stimulation for FOS induction**

Animals were lightly restrained using a rodent restraint bag and one of the hind paws was stroked lightly with a #5 painting brush. Each 2-second stroke was applied from the middle of the foot to the distal foot pad along the peripheral side to cover the sural nerve territory. These strokes were applied once every 4 s for 10 min[11]. This touch stimulus does not elicit a flexion reflex in normal mice. Animals were perfused 1.5 hr later with 4% PFA and spinal cord and brain tissues were postfixed in 30% sucrose and 4% PFA for 2 additional days. 30 μm coronal sections of hole brain and cross sections of spinal cord L3-L5 were obtained and stored in PBS at 4 °C. After permeabilization and blocking, slices were subsequently incubated with a FOS antibody (ABE457, EMD Millipore) overnight at 4 °C, followed by incubation with an Alexa Fluor 546 donkey anti-rabbit secondary anti-body (ThermoFisher, cat#A10040) for 1 h at room temperature. The slices were then washed and mounted on slides with Vectashield-containing DAPI (H-1500, Vector Laboratories). Images were taken from 2 or 3 slices per animal. A minimal of 3 animals were analyzed for each group. The number of FOSh+ cells was quantified in a fixed area within sections.

**Electrophysiology**

Acute spinal cord slice preparation: 5–7-week-old mice were anesthetized with isoflurane, decapitated, and the vertebral column was rapidly removed and placed in oxygenated ice-cold dissection solution containing (in mM): 250 sucrose, 2.5 KCl, 25 $NaHCO_3$, 1 $NaH_2PO_4$, 25 glucose, 6 $MgCl_2$, 0.5 $CaCl_2$, and 5 kynurenic acid (pH = 7.4, 320 mOsm). The lumbar spinal cord was isolated, embedded in a 3% low melting point agarose block and transverse of parasagittal slices (250 μm thick) with dorsal roots attached were made using a vibrating microtome (Leica VT1200). Slices were incubated for 10-15 min in oxygenated recovery solution containing (in mM): 92 NMDG, 2.5 KCl, 1.2 $NaH_2PO_4$, 30 $NaHCO_3$, HEPES 20, Glucose 25, 5 Sodium Ascorbate, 2 thiourea, 3 Sodium Pyruvate, 10 $MgSO_4$, 0.5 $CaCl_2$ (pH = 7.3~7.4, 300~310 mOsm) then transferred to aCSF containing (in mM): 125 NaCl, 2.5 KCl, 1.0 $NaH_2PO_4$, 25 $NaHCO_3$, 25 glucose, 1 $MgCl_2$, and 2 $CaCl_2$ (pH = 7.4, 320 mOsm). For Acute brain slices preparation: Following isoflurane anesthesia, mice were decapitated and brains were quickly removed and transferred into ice-cold high sucrose aCSF. Coronal slices of 300 μm were made and allowed to recover at 32 °C – 34 °C for 30 min before being moved to ACSF at room temperature[5].

Patch-clamp recordings of PV+ neurons were performed at room temperature. PV+ neurons located within or closely juxtaposed to the substantia gelatinosa in the spinal cord, or within layer II/III of the anterior cingulate cortex were identified with a Cre-dependent YFP reporter. For spinal cord recordings, glass pipettes (5-6 MΩ tip resistance) were filled with an internal solution containing (in mM): 120 K-methyl-sulfonate, 10 NaCl, 10 EGTA, 1 $CaCl_2$, 10 HEPES, 0.5 $Na_3GTP$, 5 MgATP (pH = 7.3, 310-320 mOsm). For cortical recordings, recording pipettes (3-4 MΩ) were filled with an internal solution containing (in mM): 130 K-gluconate, 10 KCl, 10 HEPES, 5 MgATP, 0.3 $Na_3GTP$, and 0.2 EGTA (pH = 7.3, 300~310 mOsm). Membrane excitability experiments were performed in current-clamp mode with injections of incremental DC current steps (800 ms, -50 to +250 pA with a step interval of 20 pA). The numbers of action potentials elicited by injected currents were counted. Data were acquired using a Multiclamp 700B amplifier and pClamp10 software (Molecular Devices, USA). Sampling rate was 10 kHz.

For synaptic inhibition experiments, a floxed ChR2 line (stock # 012569) was crossed with the PV-Cre driver line to allow optogenetic activation of PV+ terminals. Patch-clamp recordings were targeted in PV/ChR2-negative neurons in substantia gelatinosa for both presynaptic inhibition and postsynaptic inhibition assessment. Recording pipettes (3-4 MΩ) were filled with an internal solution containing (in mM): 132 $CsMeSO_3$, 8 CsCl, 10 HEPES, 0.6 EGTA, 4 MgATP, 0.4 $Na_3GTP$, and 10 Na-phosphocreatine (pH = 7.3, 300~310 mOsm). Full-field photostimulation (PS) of PV-ChR2-expressing neurons was achieved using single light pulses to induce maximal responses (450nm wavelength, 1 ms, 30 mW, Laser Diode MDL-III-450), which was collimated and coupled to the epifluorescence path of an Olympus BX51 microscope. For presynaptic inhibition through axo-axonic synapse onto primary afferent terminals, primary afferent depolarization (PAD)-induced optically evoked EPSC was recorded in postsynaptic neurons at the reversal potential of IPSCs (-70 mV) at room temperature[12]. For postsynaptic inhibition, glutamate receptor antagonists CNQX (10 µM) and APV (100 µM) were added to the recording aCSF, and optically evoked IPSCs were recorded in the postsynaptic neurons held at 0 mV.

**Behavioral tests**

**Open field test:** Mice were placed in a 40 cm (L) x 40 cm (W) x 40 cm (H) open-field chamber. Locomotor activity was recorded for 30 minutes using an overhead digital camera and tracked using Viewer III tracking system (BIOSERVE, Bonn, Germany). Time spent in the center was measured by a 10 cm by 10 cm area in the center.

**Elevated plus maze:** An elevated plus maze (Stoelting Co. IL) is composed of two open arms and two closed arms that extend from a central square area and elevate to a height of 50cm above floor level. The mouse was placed in the center of the maze facing an open arm and left to freely explore the maze for 10 min. The amounts of the time spent in the open vs. closed arms were recorded by Viewer III tracking system (BIOBSERVE).

**Y-Maze test:** A plastic y-maze (Stoelting Co. IL) was used to measure spatial working memory. Individual mice were placed in the center of the Y-maze and allowed to freely explore for 5 minutes. The sequences and total numbers of arm entries were recorded and analyzed with the Viewer III tracking system. Visiting all three different arms consecutively was termed a 'correct' trial and visiting one arm twice or more than three consecutive entries was termed a 'wrong' trial. Spontaneous alternation was calculated as the percentage of the 'correct' trial to the total trials.

**Hotplate test:** Hot plate test was performed using a hot plate with custom floorless Plexiglas chamber (TAP Plastic, Mountain View, CA). Mice were acclimated to the testing environment for 20 minutes. The plate temperature was set to 50 °C and 55 °C. Mice were placed on the plate and the latency to withdraw/lick a hind paw was scored. A cut-off of 60 s was set to prevent tissue damage.

**Formalin test:** Mice received a single subcutaneous injection of 50 µl of 1% formalin in the plantar surface of a single hind paw delivered with a 26-G needle. Spontaneous pain behavior, characterized by increased paw flinching, licking, and elevated paw were scored in the 60 min period after the injection.

**Von Frey withdrawal threshold test:** Each mouse was habituated in a small (7.5 × 7.5 × 15 cm) plastic cage for at least 20 minutes before testing. Mechanical sensitivity was determined with a series of von Frey filaments (bending forces: 0.04, 0.07, 0.16, 0.4, 0.6, 1, 1.4, 2 and 4 g) applied within the sciatic nerve territory (lateral part of the hind paw) to assess mechanical withdrawal thresholds. Filaments were applied perpendicular to the hind paw surface with sufficient force to cause a slight bending of the filament. A positive response was characterized by a rapid withdrawal of the paw away from the stimulus fiber within 4 s. The Up-Down method was used to determine the mechanical threshold (50% withdrawal threshold). The 50% threshold was calculated using the formula: 50% threshold (g) = $10^{(X+kd)}/10^4$, where X = the value (in log units) of the final von Frey filament, k = tabular value for the response pattern[13,14] and d = the average increment (in log units between von Frey filaments).

**Statistics**

All experiments were conducted with the experimenters being "blind" to the genotypes and treatment parameters. Animals in the same litter were randomly assigned to different treatment groups and blinded to experimenters. Sample sizes (n number of independent biological replicates) were first determined based on similar experiments performed and published by our lab and others in the same field[4,11,15]. To ensure reproducibility, each experiment was performed in at least three independent litters (N number), with multiple animals per litter. For non-behavioral experiments (e.g. histology), a minimum of 3 technical duplicates were analyzed and results averaged to generate a single data point for that particular animal. All samples in each group were analyzed without exclusions with one exception: if the injection site was found inaccurate during end-point histological verification, all data from that particular animal were excluded. The n/N (number of cells/number of mice) for each experiment are indicated in figure legends.

All results are presented as mean ± SEM, and statistical analysis were performed using GraphPad Prism 7 software (GraphPad Software, San Diego, CA). The distribution of data in each set of experiments was tested for normality using Shapiro-Wilk normality test. Two-tailed unpaired $t$ test (for parametric test) or Mann-Whitney $U$ tests (for nonparametric test) was applied for two groups comparisons. For multiple groups comparisons, one-way ANOVA (for parametric test) or Kruskal-Wallis test (for nonparametric test) were applied, followed by appropriate *post hoc* tests (as indicated in figure legends).

### References


1 Chapellier, B. *et al.* A conditional floxed (loxP-flanked) allele for the retinoic acid receptor alpha (RARalpha) gene. *Genesis* **32**, 87-90 (2002).
2 Sarti, F., Schroeder, J., Aoto, J. & Chen, L. Conditional RARalpha knockout mice reveal acute requirement for retinoic acid and RARalpha in homeostatic plasticity. *Front Mol Neurosci* **5**, 16, doi:10.3389/fnmol.2012.00016 (2012).
3 Yee, A. X. & Chen, L. Differential regulation of spontaneous and evoked inhibitory synaptic transmission in somatosensory cortex by retinoic acid. *Synapse* **70**, 445-452, doi:10.1002/syn.21921 (2016).
4 Wang, D. *et al.* Functional Divergence of Delta and Mu Opioid Receptor Organization in CNS Pain Circuits. *Neuron* **98**, 90-+, doi:10.1016/j.neuron.2018.03.002 (2018).
5 Zhong, L. R., Chen, X., Park, E., Sudhof, T. C. & Chen, X. Retinoic Acid Receptor RAR alpha-Dependent Synaptic Signaling Mediates Homeostatic Synaptic Plasticity at the Inhibitory Synapses of Mouse Visual Cortex. *Journal of Neuroscience* **38**, 10454-10466, doi:10.1523/Jneurosci.1133-18.2018 (2018).
6 Decosterd, I. & Woolf, C. J. Spared nerve injury: an animal model of persistent peripheral neuropathic pain. *Pain* **87**, 149-158, doi:Doi 10.1016/S0304-3959(00)00276-1 (2000).
7 Zolotukhin, S. *et al.* Recombinant adeno-associated virus purification using novel methods improves infectious titer and yield. *Gene Ther* **6**, 973-985, doi:10.1038/sj.gt.3300938 (1999).
8 Peirs, C. *et al.* Dorsal Horn Circuits for Persistent Mechanical Pain. *Neuron* **87**, 797-812, doi:10.1016/j.neuron.2015.07.029 (2015).
9 Cetin, A., Komai, S., Eliava, M., Seeburg, P. H. & Osten, P. Stereotaxic gene delivery in the rodent brain. *Nat Protoc* **1**, 3166-3173, doi:10.1038/nprot.2006.450 (2006).
10 Park, E., Tjia, M., Zuo, Y. & Chen, L. Postnatal Ablation of Synaptic Retinoic Acid Signaling Impairs Cortical Information Processing and Sensory Discrimination in Mice. *J Neurosci* **38**, 5277-5288, doi:10.1523/JNEUROSCI.3028-17.2018 (2018).
11 Liu, Y. *et al.* Touch and tactile neuropathic pain sensitivity are set by corticospinal projections. *Nature* **561**, 547-550, doi:10.1038/s41586-018-0515-2 (2018).
12 Boyle, K. A. *et al.* Defining a Spinal Microcircuit that Gates Myelinated Afferent Input: Implications for Tactile Allodynia. *Cell Rep* **28**, 526-+, doi:10.1016/j.celrep.2019.06.040 (2019).
13 Chaplan, S. R., Bach, F. W., Pogrel, J. W., Chung, J. M. & Yaksh, T. L. Quantitative assessment of tactile allodynia in the rat paw. *J Neurosci Methods* **53**, 55-63, doi:10.1016/0165-0270(94)90144-9 (1994).
14 Gonzalez-Cano, R. *et al.* Up-Down Reader: An Open Source Program for Efficiently Processing 50% von Frey Thresholds. *Frontiers in Pharmacology* **9** (2018).
15 Hsu, Y. T., Li, J., Wu, D., Sudhof, T. C. & Chen, L. Synaptic retinoic acid receptor signaling mediates mTOR-dependent metaplasticity that controls hippocampal learning. *Proc Natl Acad Sci U S A* **116**, 7113-7122, doi:10.1073/pnas.1820690116 (2019).

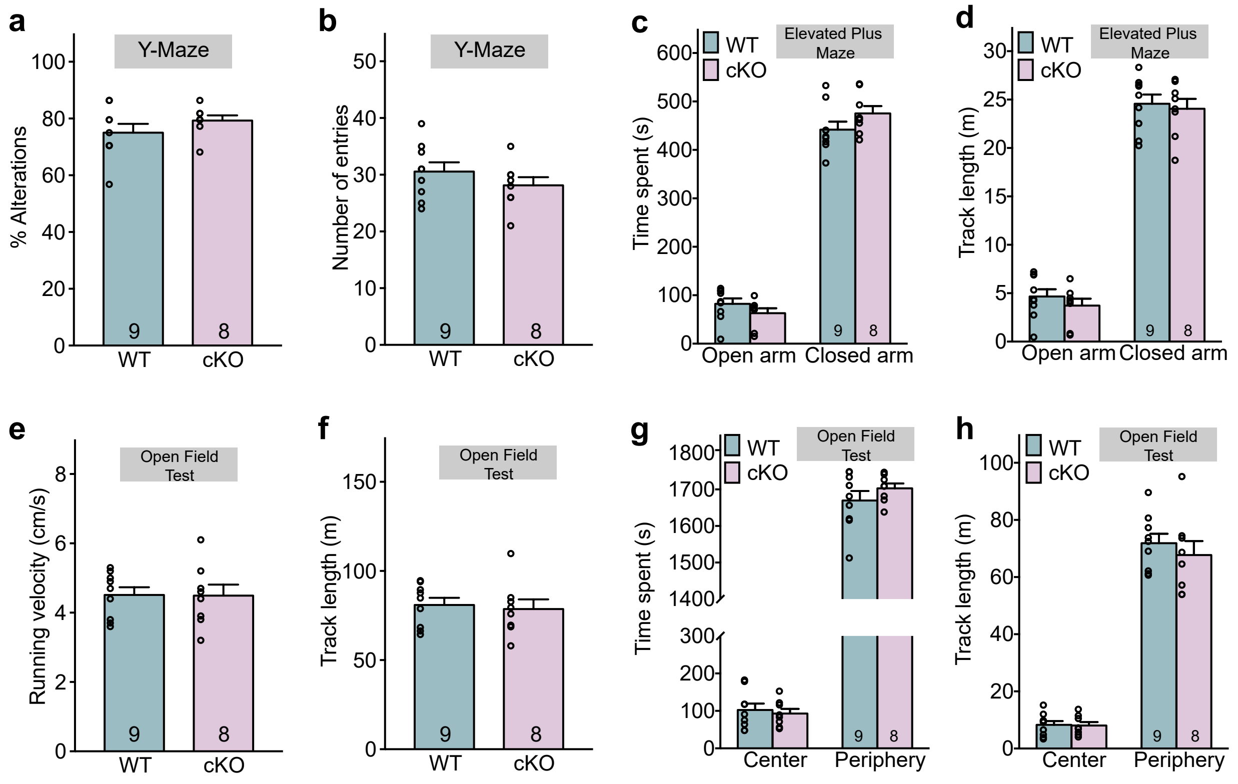


**Extended Data Fig. 1 | Behavioral assessment of Y-Maze, elevated plus maze and open field test in wild-type and PV-RARα conditional knock-out mice.**

**a-b.** Quantification of Y-maze test. Percentage of alternations (**a**) and number of total entries (**b**) were measured from WT and PV-RARα KO mice.

**c-d.** Quantification of Elevated plus maze test. Average time spent (**c**) and average track length in open or closed arms (**d**) were measured.

**e-h.** Quantification of open field test. Aaverage velocity (**e**), average track length (**f**), average time spend (**g**), and average track length (**h**) in the center and periphery of the chamber were measured. All data represent mean ± SEM.

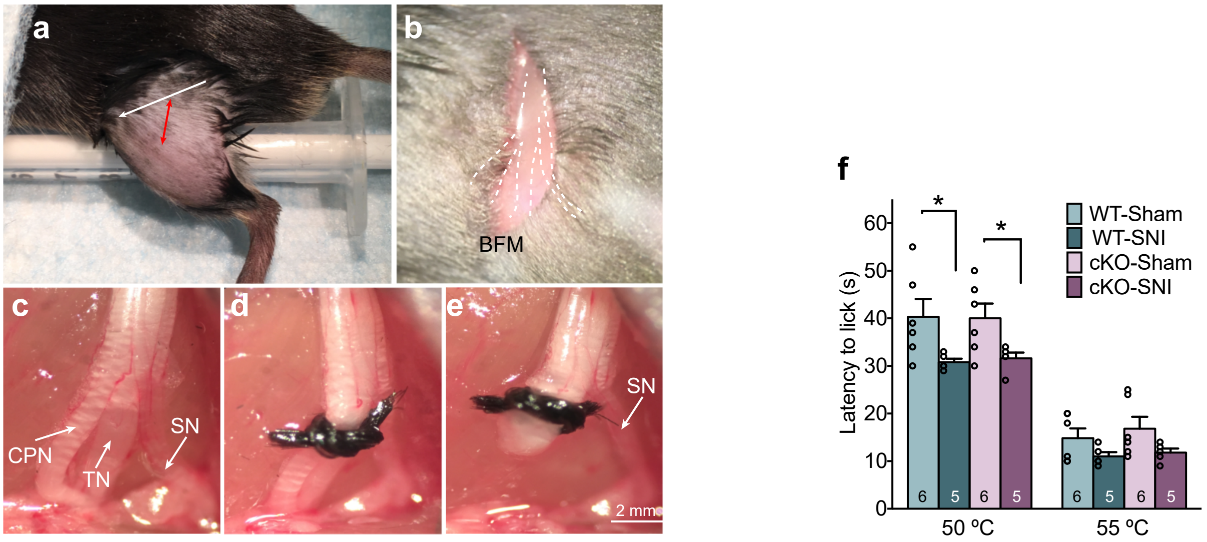


**Extended Data Fig. 2 | SNI surgery and quantification of hot plate test in WT and PV-RARα cKO mice**.

**a.** The incision (red line) is made at the middle position of the thigh at a 60º angle from a fictive axe (white line) between trochanter major and iliaca cresta.
**b.** Following the incision along the red line, expose the biceps femoris muscle (BFM) and perform a careful blunt dissection to expose the trifurcation of the sciatic nerve. Dotted lines show the position of the sciatic trifurcation underneath the artery genus descendes and biceps femoris.
**c.** The three bare branches of the sciatic nerve are shown. common peroneal (CPN), tibial (TN) and sural nerves (SN).
**d.** Ligation of the common peroneal and tibial nerves was performed with a surgical knot.
**e.** The ligated nerves were transected distally for 2 mm section to prevent nerve regeneration. Care was taken to avoid contact with the sural nerve.
**f**. Thermal hyperalgesia quantified as latency to licking hind paws when placed onto 50 $^{\circ}$C and 55 $^{\circ}$C hot plate in WT. *, $p < 0.05$; two-sided unpaired t-test. Data shown as mean ± s.e.m..

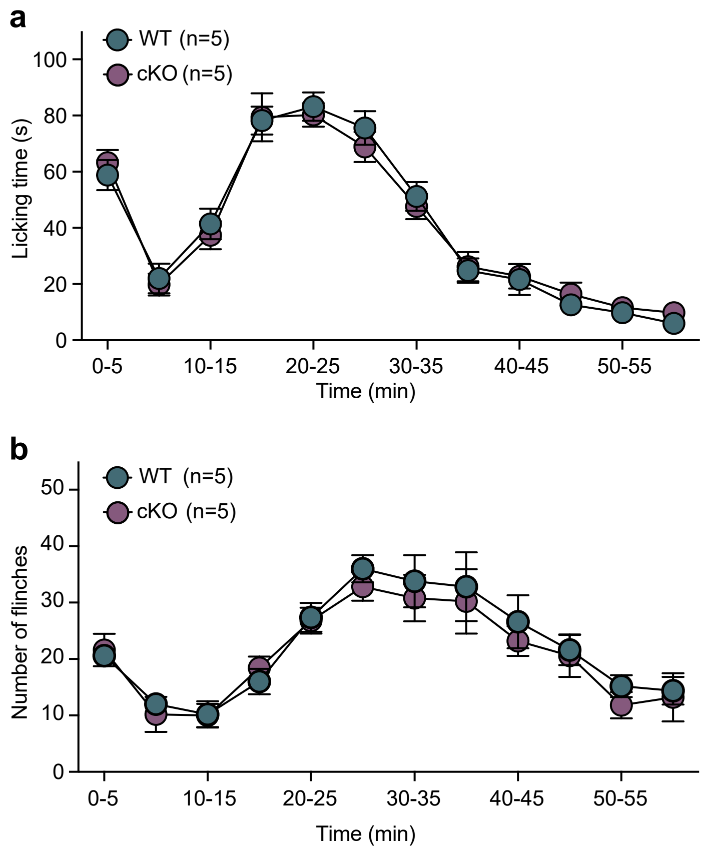


**Extended Data Fig. 3 | Assess nociceptive behavior through formalin test.**
**a-b.** Time course of paw licking responses and flinching in WT and PV-RARα cKO mice following injection of 1% formalin (50µl, subcutaneous) under the plantar of left hind paw at T=0. Both WT and cKO mice showed the two characteristic phases of nociceptive behavior: An early phase of high licking/flinch period immediately after formalin injection that lasted approximately 5 min, followed by a lull, and a more prolonged but delayed late phase. Behavioral measurements are binned every 5 minutes (mean ± s.e.m.).

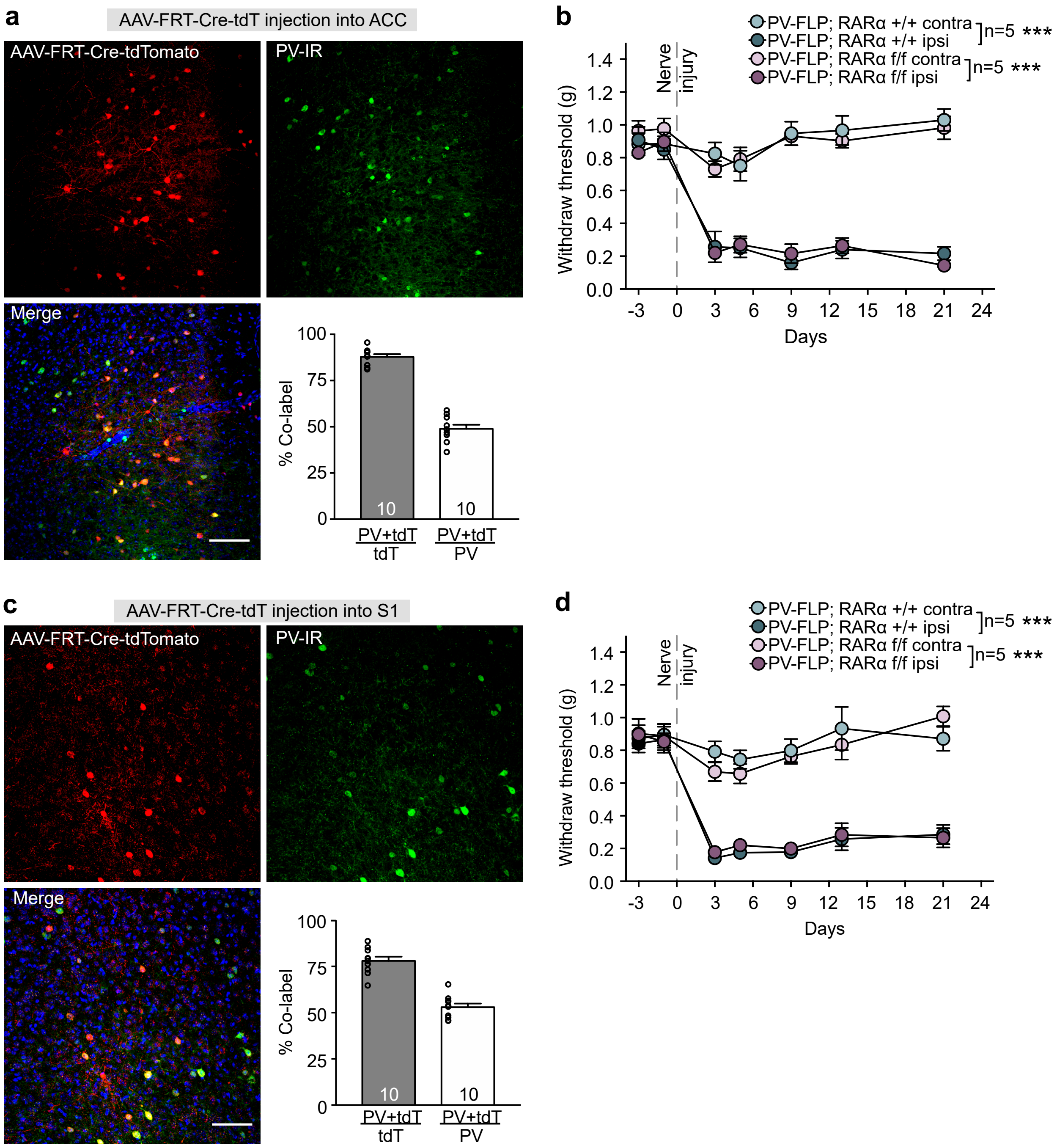


**Extended Data Fig. 4 | Region-specifically deletion of RARα in the ACC or S1 does not block SNI-induced mechanical allodynia.**

**a.** Representative images showing FRT-Cre-tdTomato expression in PV-immunoreactive ACC neurons of PV-Flp driver line crossed with floxed RARα cKO mice. Scale bar: 100 µm. Bar graph shows PV-Flp driver line expression fidelity (% PV-IR/tdTomato) and efficacy (% tdTomato/PV-IR) in ACC.

**b.** Mechanical allodynia quantified as withdraw threshold in von Frey test in WT and ACC-specific PV-RARα KO mice. ***, $p < 0.001$; two-way ANOVA followed by Bonferroni test.

**c.** Representative images showing FRT-Cre-tdTomato expression in PV-immunoreactive S1 neurons of PV-Flp driver line crossed with floxed RARα cKO mice. Scale bar: 100 µm. Bar graph shows PV-Flp driver line expression fidelity (% PV-IR/tdTomato) and efficacy (% tdTomato/PV-IR) in S1.

**d.** Mechanical allodynia quantified as withdraw threshold in von Frey test in WT and S1-specific PV-RARα KO mice. ***, $p < 0.001$; two-way ANOVA followed by Bonferroni test. All N = # of mice. All data shown as mean ± s.e.m..

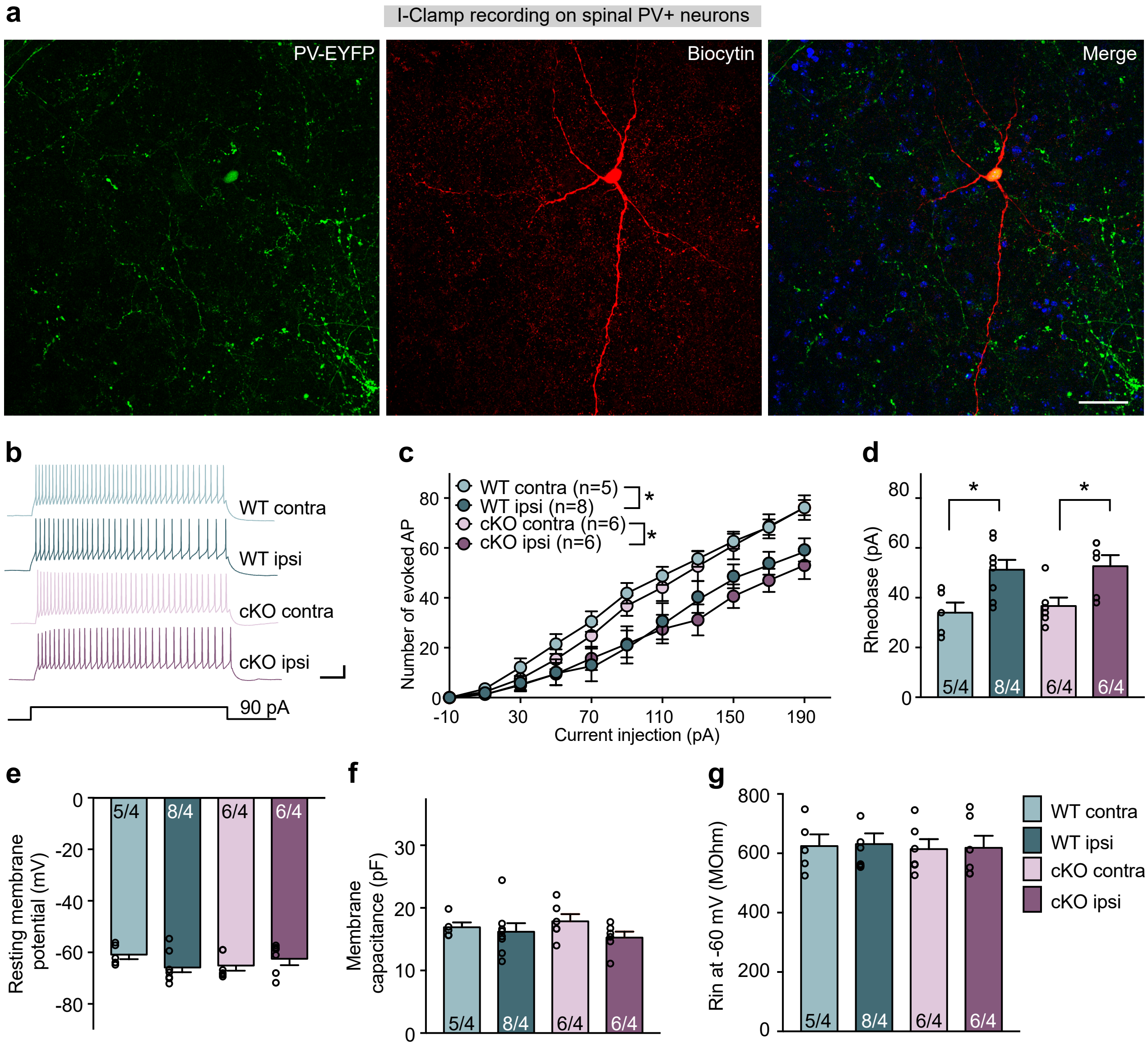


**Extended Data Fig. 5 | Spared nerve injury affects active membrane properties of PV+ cells in the dorsal horn.**

**a.** Representative images of biocytin-labelled (red) PV+ cells, which was identified with Cre-dependent EYFP reporter expression.

**b.** Sample traces of current clamp recordings from WT and RARα KO PV+ neurons ipsi- and contralateral to SNI side in response to a step current injection of 90 pA. Scale bars: 100 ms, 10 mV.

**c**. Input-output relationship between the total number of APs and the step current injections. *, $p < 0.05$, two-way ANOVA followed by Bonferroni test.

**d.** Rheobase of PV+ cells at the ipsilateral and contralateral sides of WT and cKO mice.
*, $p < 0.05$, two-way ANOVA followed by Bonferroni test.

**e-g.** Quantification of passive membrane properties measured as resting membrane potentials (**e**), membrane capacitances (**f**), and input resistances (**g**) at -60 mV holding potential.
All data shown as mean ± s.e.m.

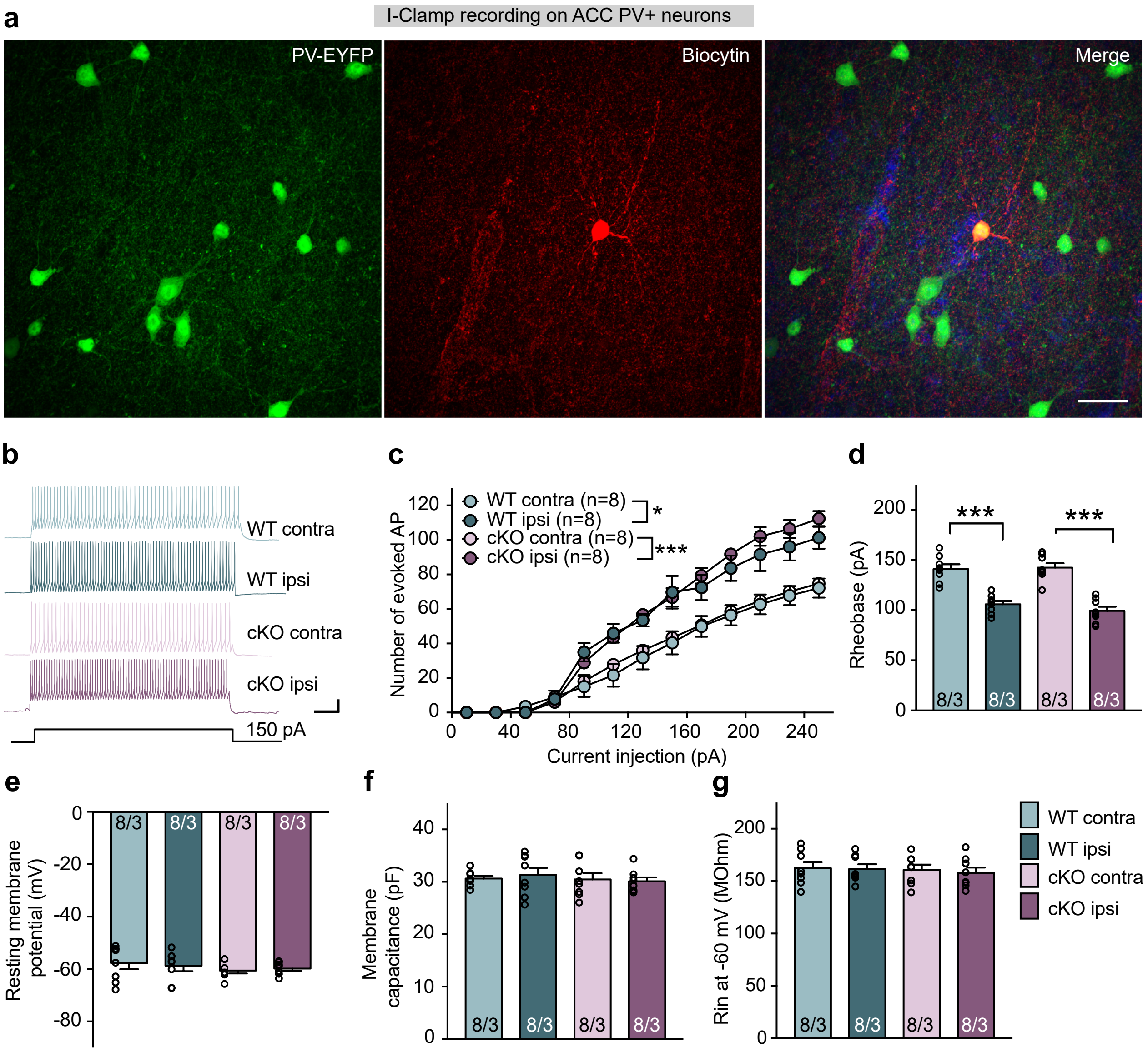


**Extended Data Fig. 6 | Spared nerve injury affects active membrane properties of PV+ cells in the ACC.**

**a.** Representative images of biocytin-labelled (red) PV+ cells, which was identified with Cre-dependent EYFP reporter expression.

**b.** Sample traces of current clamp recordings from WT and RARα KO PV+ neurons ipsi- and contralateral to SNI side in response to a step current injection of 150 pA. Scale bars: 100 ms, 10 mV.

**c**. Input-output relationship between the total number of APs and the step current injections. *, $p < 0.05$; ***, p < 0.001; two-way ANOVA followed by Bonferroni test.

**d.** Rheobase of PV+ cells at the ipsilateral and contralateral sides of WT and cKO mice.
***, $p < 0.001$, two-way ANOVA followed by Bonferroni test.

**e-g.** Quantification of passive membrane properties measured as resting membrane potentials (**e**), membrane capacitances (**f**), and input resistances (**g**) at -60 mV holding potential.
All data shown as mean ± s.e.m.

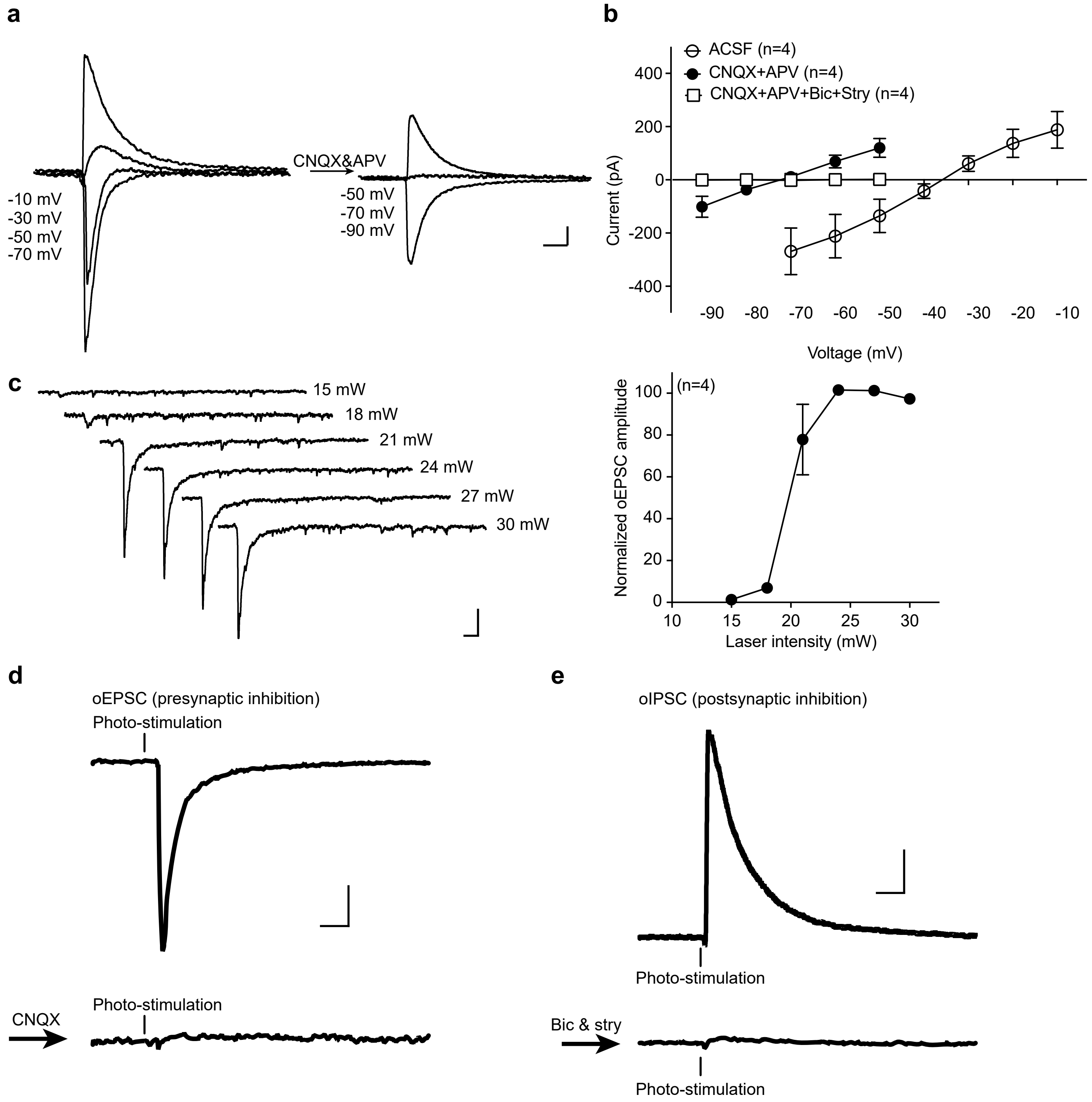


**Extended Data Fig. 7** | Pharmacologically characterization of postsynaptic responses in PV+ target neurons.

**a.** Example traces of oPSCs at different membrane holding potentials in response to photo-stimulation of ChR2-expressing PV+ cells in ACSF (left) and after adding CNQX (10 μM) and APV (100 μM, right). Scale bars: 50 ms, 20 pA.

**b.** I-V curve of oPSCs in ACSF, CNQX + APV and CNQX + APV + bicuculine + strychnine. In the presence of CNQX + APV, oIPSCs exibit a reversal potential of -70 mV, and can be further blocked by bicuculine and strychnine.

**c.** Representative traces (left) and input-output curve (right) of oEPSCs from postsynaptic neurons receiving PV+ neuron inputs. Scale bars: 50 ms, 20 pA.

**d.** oEPSCs evoked by photo-stimulation of PV+ neurons can be completely blocked by AMPA receptor antagonist CNQX. Scale bars: 50 ms, 20 pA.

**e.** oIPSCs evoked by photo-stimulation PV+ neurons can be completely blocked by GABA and glycine receptor antagonist bicuculline (10 μM) and strychnine (1 μM). Scale bars: 50 ms, 20 pA.
